# *phylogatR*: Phylogeographic data aggregation and repurposing

**DOI:** 10.1101/2021.10.11.461680

**Authors:** Tara A Pelletier, Danielle J Parsons, Sydney K Decker, Stephanie Crouch, Eric Franz, Jeffery Ohrstrom, Bryan C Carstens

## Abstract

Patterns of genetic diversity within species contain information about the history of that species, including how they have responded to historical climate change and how easily the organism is able to disperse across its habitat. More than 40,000 phylogeographic and population genetic investigations have been published to date, each collecting genetic data from hundreds of samples. Despite these millions of data points, meta-analyses are challenging because the synthesis of results across hundreds of studies, each using different methods and forms of analysis, is a daunting and time-consuming task. It is more efficient to proceed by repurposing existing data and using automated data analysis. To facilitate data repurposing, we created a database (*phylogatR)* that aggregates data from different sources and conducts automated multiple sequence alignments and data curation to provide users with nearly ready-to-analyze sets of data for thousands of species. Two types of scientific research will be made easier by *phylogatR*, large meta-analyses of thousands of species that can address classic questions in evolutionary biology and ecology and student- or citizen-science based investigations that will introduce a broad range of people to the analysis of genetic data. *phylogatR* enhances the value of existing data via the creation of software and web-based tools that enable these data to be recycled and reanalyzed and increase accessibility to big data for research labs and classroom instructors with limited computational expertise and resources.

## Introduction

Quantifying the geographic distribution of genetic variation within and between species provides essential information for understanding the evolutionary processes that give rise to current biodiversity patterns and is an essential aim of landscape genetic and phylogeographic investigations. The NCBI GenBank database houses over two hundred million DNA sequences, a number that grows monthly (https://www.ncbi.nlm.nih.gov/genbank/statistics/), but most of these sequences lack metadata associated with the locality from which the organism was collected. This limits the potential use of these data by preventing repurposing of the data (e.g., Sidlauskas et al. 2010) in any analyses that require geospatial information. For example, Marques et al. (2013) found that only 7% of GenBank accessions of barcoding genes, such as *Cytochrome Oxidase I* (*COI*), include latitude and longitude, and only 18% list museum catalogue information that can be used to link the sequence to a particular specimen. Similarly, Gratton et al. (2017) found that only 6.2% of GenBank tetrapod accessions include locality data. Overall, it has been suggested that 90% of biodiversity data remain unavailable for further use, and that missing geographic information was the most significant factor limiting use (Peterson et al. 2018). These ‘missing’ locality data are particularly problematic when it is understood that voucher specimens from thousands of investigations are deposited into natural history collections, and that metadata associated with these vouchers, including in many cases georeferenced locality data, are currently available in other databases such as the Global Biodiversity Information Facility (GBIF).

Spatial information is extremely important to the biological sciences. For example, more than 22,000 published papers use some variant of the word “phylogeo*” in their title or abstract, in addition to more than 22,000 that use “population genetics” (https://www.webofscience.com/wos/woscc/basic-search, 9-Sept-2021). These disciplines necessarily include spatial information, and this component enables researchers to explore topics such as speciation (e.g., Smith & Carstens 2020), hybridization (Burbrink et al. 2021), demographic change (e.g., Carstens et al. 2018), and estimating the current (e.g., Farallo et al. 2020), former (e.g., Pelletier & Carstens 2016) or future (e.g., Nottingham & Pelletier 2021) species ranges, in addition to the evaluation of ecological niche overlap (e.g., Cavalcante et al. 2020). Given that researchers in each of these disciplines routinely collect sequence data from hundreds of samples (Garrick et al. 2015), the existence of georeferenced data in databases such as GenBank and BOLD (Barcode of Life Database) can enable novel comparative analyses.

Large-scale meta-analyses offer a promising strategy to understand the broad-scale effects of geography, geology, and climate change on species distributions (Guralnick & Hill 2009) and hold immense potential for insight (Dawson 2014; Heberling et al. 2021). However, the considerable variation in study design and statistical analyses used across studies render meta-analysis in population genetics and phylogeography difficult (Garrick et al. 2015). A more productive strategy is the repurposing of data (Sidlauskas et al. 2010; Blanchet et al. 2017; Leigh et al. 2021), where data from previously published work are reanalyzed in large groups in order to extract insight about global processes. Combining similar types of data from multiple studies and then re-analyzing these data under a common framework has the power to elucidate factors that drive evolution on both small and large scales.

One example of the potential of data repurposing is found in Miraldo et al. (2016). These researchers manually assembled mitochondrial DNA (mtDNA) sequences from almost 2000 species of terrestrial mammals and amphibians and used these data to document that genetic diversity is higher in the tropics and lower where human populations are high. This analysis required a considerable amount of effort, as data were mined by downloading GenBank and BOLD accessions that contained geographic coordinates or by emailing researchers to ask for their data. The data curation in Miraldo et al. (2016) was manual, which places an upper limit on the number of species that can be included in the analysis. More recent investigations have used automated computational pipelines to increase the efficiency of exploring population genetics and species limits on large scales in several ways. For example, Pelletier and Carstens (2018) used a Python script to assemble a database of over 8000 species of plants, fungi, and animals, analyzed these data using R, and demonstrated that genetic structure within species was higher in northern latitudes and that the size of a species range was an important predictor of genetic structure.

Existing macrogenetic studies demonstrate the need for global analyses of genetic data. Large-scale biodiversity data enhances conservation efforts (Pelletier et al. 2018; Thompson et al. 2021) and mapping the tree of life (Folk & Siniscalchi 2021). There is a strong push for making data publicly available (Marden et al. 2021) and repurposing these data increases their value (Whitlock et al. 2010; Heberling et al. 2021). It opens the doors for reexamining classic questions on larger scales, but also moves forward the fields of population genetics, phylogeography, and systematics by increasing the power to tease apart the complex processes that shape biodiversity patterns (Hickerson et al. 2010; Papadopoulou & Knowles 2016). Furthermore, integrating data types (e.g., environmental data layers, morphological measurements, life history characteristics) with large-scale genetic and geographic data will not only enhance our understanding of the ecological processes that contribute to evolutionary change, but also provide applicable information for conservation purposes (Anderson et al. 2020).

In order to facilitate phylogeographic analyses on the largest possible scale (i.e., continental or global) from thousands of species, we have developed software that parses accessions from several repositories of geographic and genetic information, organizes them into a common framework under a taxonomic hierarchy, and produces multiple sequence alignments that are analysis-ready. Our goal was to develop a database that is user-friendly and accessible to researchers and instructors without training in computational biology whose efforts are aimed at conducting studies on specific taxonomic groups and/or biogeographic regions. This effort contributes to FAIR (Findability, Accessibility, Interoperability, and Reusability) initiatives that aim to improve the infrastructure of open-data science (Wilkinson et al. 2016; Heberling et al. 2021). The database, *phylogatR* (<u>*phylo*</u>geographic data aggre<u>*gat*</u>ion and <u>*r*</u>epurposing) is freely available via the Ohio Supercomputer Center (OSC), along with several <u>R</u> scripts to aid in data curation, analysis, and education.

## *phylogatR* pipeline

Data for our aggregator comes primarily from three large databases. 1) GBIF (https://www.gbif.org/), an open-source database funded and supported since 1999 by a large group of government agencies worldwide. It contains over 1 billion occurrence records from over 6 million organisms across the globe. 2) NCBI GenBank, a collection of DNA sequence data from three organizations: DNA DataBank of Japan (DDBJ), the European Nucleotide Archive (ENA), and GenBank at NCBI, (https://www.ncbi.nlm.nih.gov/genbank/), and 3) BOLD http://www.boldsystems.org/index.php), developed by the Center for Biodiversity Genomics in Canada, contains barcode data for almost 600,000 species. Pipeline choices were made to minimize data duplication and loss, conduct preliminary cleaning and alignment, and to return results to users in a manner that is transparent and enables them to conduct additional curation as needed. Scripts for data aggregation and cleaning are available in our GitHub repository (https://github.com/OSC/phylogatr-web). A schematic overview of the pipeline is available in Figure 1.

**Figure 1.**
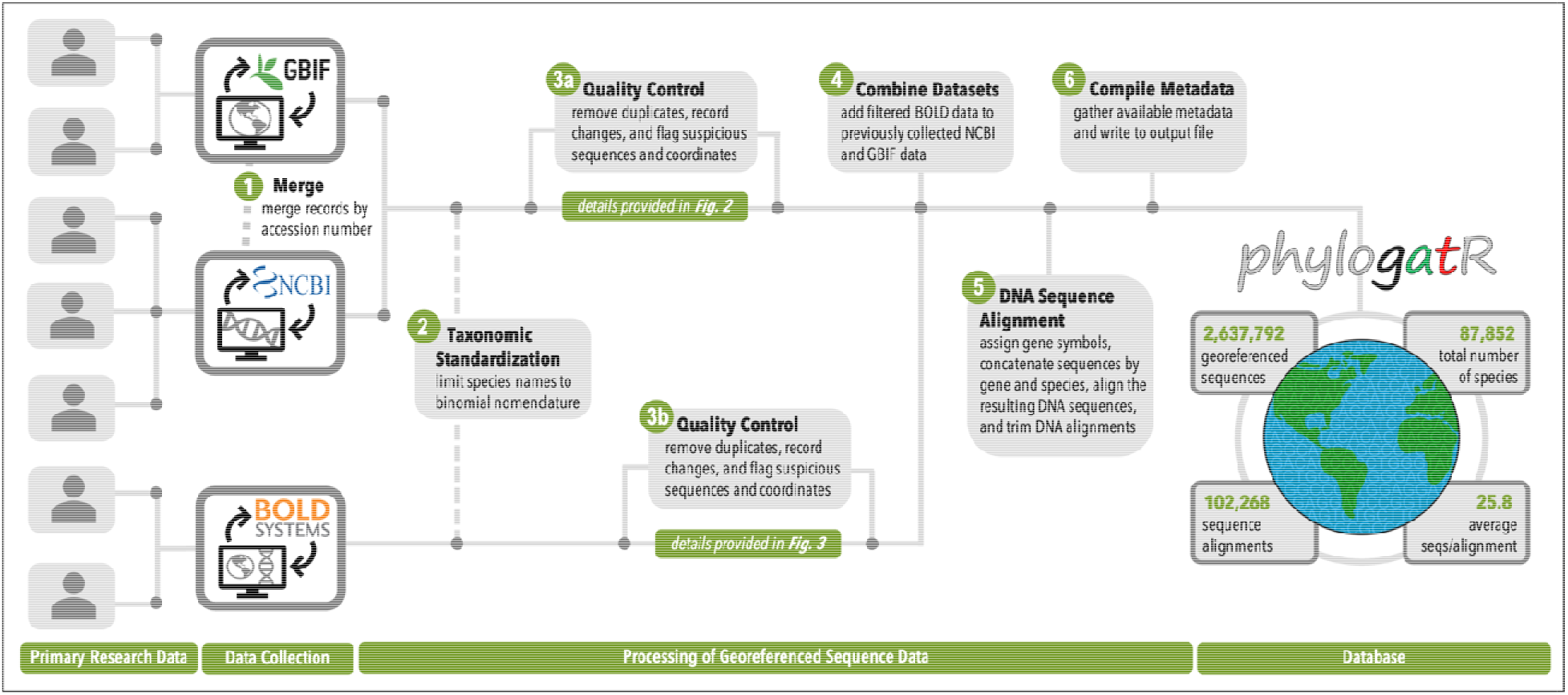
Overview of *phylogatR* pipeline.

### Data aggregation

Data were downloaded from GBIF that included coordinates, excluding those flagged as suspicious, contained sequence accessions, and a full binomial name. We only included occurrences in which Basis of record was either PreservedSpecimen, MaterialSample, HumanObservation, or MachineObservation. The entire GenBank nucleotide sequence database was downloaded using the rsync file transfer program. Occurrences and DNA sequences that contained the same GenBank accession were matched and curated (Fig. 2). For each occurrence, sequence accessions and geographic coordinates were checked for duplication. First, all coordinates were rounded to two decimals to overcome differences in coordinates that come from the same sample but appear different due to rounding. If coordinates were different, but had the same GenBank accession, we assumed duplicates represent different individuals uploaded to GenBank as a single haplotype. In this case, all occurrences were kept, but each was flagged with ‘**g**’ so that users can explore these accessions if necessary. If coordinates were the same, we checked the basis of record. If these were different, we kept only the highest precedence for an observation (from high to low: preserved specimen, material sample, human observation, machine observation), with the assumption that these sequences with the same GenBank accession and geographic coordinates was a different observation of the same specimen, and each was flagged with ‘**b**’. If basis of record was the same, we checked the species name. If different, we assumed a change in taxonomy and kept the most recent occurrence and flagged it with ‘**s**’. If the species name was also the same, we checked the event date. If different, we assume the duplicates represent different individuals, and they were flagged with ‘**d**’, again to allow further investigation by users. For any duplicates that had the same GenBank accession, geographic coordinates, species name, and event date, but different GBIF occurrences, we retained only the most recent occurrence and flagged with ‘**m**’. Next, the BOLD database was scraped to obtain taxon names and data were pulled by looping through 500 taxa at a time using the public API. All available data were downloaded and curated (Fig. 3). Records without coordinates were removed. Those with GenBank or GBIF accessions already in our database were removed.

**Figure 2.**
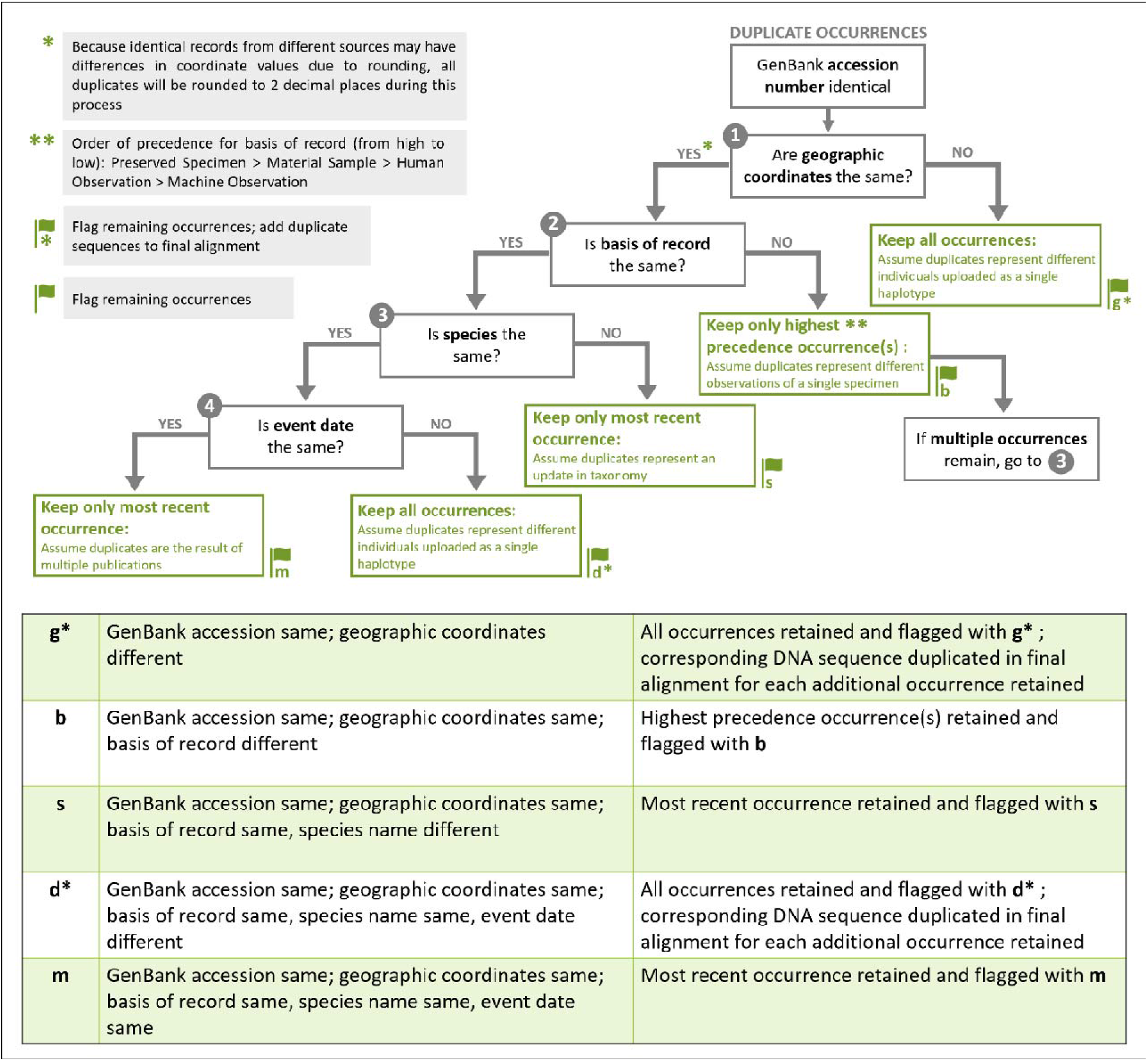
Data curation steps for GBIF and GenBank data.

**Figure 3.**
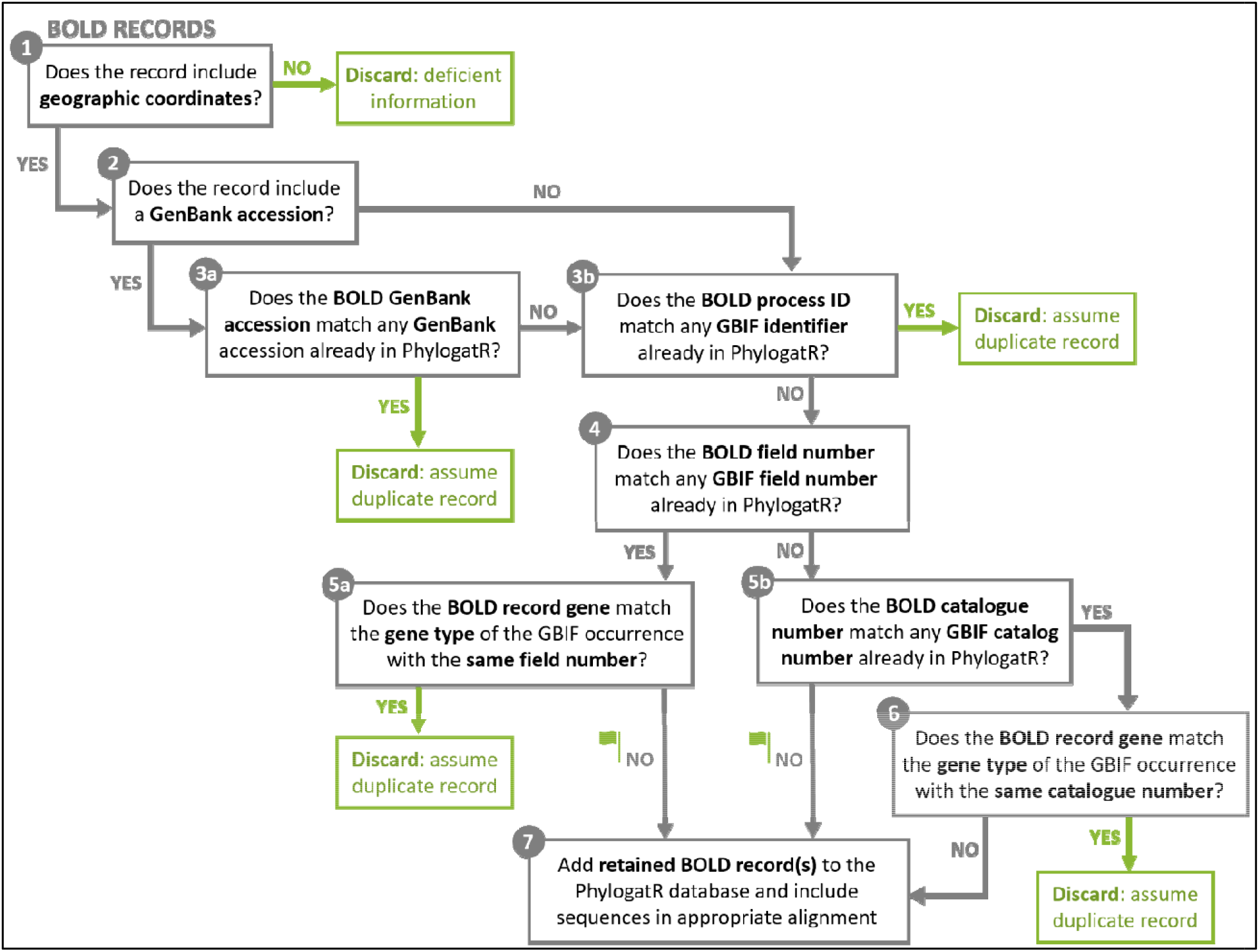
Data curation steps for BOLD data.

We standardized gene and species names to the best of our ability. For example, we assigned a common gene symbol for commonly sequenced genes that are often represented by more than one symbol (Supplemental Table S1), such as *COI* for *cytochrome oxidase I* that is also often depicted as *COXI* or *CO1*. In some cases, genes were identified with a different gene symbol but the same gene name. These were left alone assuming that they represent different regions of the same gene, such as the *malic-enzyme* that contains alignments for *ME1* and *ME2*. While we don’t expect many instances where gene symbols are incorrect, we recommend users scan the list of genes in their dataset before use. Species names were limited to binomial nomenclature, though those with subspecies identifiers are listed in the associated metadata. GBIF taxonomy was retained when it did not match the GenBank taxonomy and these are also flagged in the associated metadata. We recommend individual users to capitalize on available tools for checking taxonomy when appropriate for their needs. For example, the R package ‘taxize’ (Chamberlain & Szöcs 2013) as used in Pelletier et al. (2018) accesses many data sources to update species names, or standardized databases can be used directly to update species names such as the Mammal Diversity Database published by the American Society of Mammalogists (e.g., Parsons et al. In review).

### Multiple sequence alignment

Every sequence is identified by species, gene, GenBank accession, GBIF ID, and/or BOLD ID. All sequences were concatenated based on identical gene sequence symbol and species name. We conducted multiple sequence alignments for all genes where there were at least three sequences within a species on a species-by-species basis. First, the default MAFFT v7 parameters were used. Sequence alignments were checked by eye for 10 families (117 species-level alignments) that were previously determined to require post-alignment adjustments (Parsons et al. In review). Several alignments were found to have large sequence gaps at the ends of the alignment, while others contained unwanted sequences (e.g., parasitic sequences that have been named as the host species). After this first round of checking, only eight alignments needed trimming and three needed sequences removed (or reverse complimented). We updated the MAFFT settings to include the adjustdirection and inputorder features. Then trimAl v1.2 was used to clean the alignments. After several iterations of parameter settings, we set resoverlap to 0.85, seqoverlap to 50, and gt to 0.15. Identical sequences (same GenBank accession) with multiple GBIF occurrences that have been deemed not duplicates (Fig. 2) are repeated for the final sequence alignment. While these settings appear to eliminate most issues that arise from within species sequence alignments, researchers should screen their data for outliers before data analysis. We suspect these issues to be minimal, and when dealing with large datasets a small amount of noise is not expected to alter results (see empirical example below).

### Data

The database currently contains 87,852 species and 102,268 sequence alignments. The average number of alignments per species is 1.2 and the average number of sequences per alignment is The database includes species from Animalia (77,743), Plantae (7,905), Fungi (1,971), Chromista (229), and Protozoa (4). Most of the data are from mitochondrial and chloroplast DNA, a result which reflects the key role of genes from these organellar genomes to disciplines such as phylogeography (Garrick et al. 2015). After merging genes with different known gene symbols, our database contains a total of 1988 genes. Note that *phylogatR* has been designed to be expandable and will grow by conducting monthly scans for new accessions on GenBank, GBIF, BOLD, and other sources for data deposition for at least 10 years, and updates and fixes will be made as identified.

When data are downloaded from *phylogatR* (zip and tarball formats are available), all data are nested within directories that are structured by taxonomic rank. Each species folder consists of an unaligned fasta file (extension *.fa)* and an aligned fasta file (extension *.afa*) for each locus available for that species. Each species folder also contains an occurrence file that contains the original database accessions and geographic coordinates in decimal form, as well as any appropriate flags. The root folder contains information for each sequence alignment (in the *genes.txt* file), including the number of sequences before and after data cleaning steps, taxonomic information, and flags those that may contain inconsistencies in species names across databases. The database is available at https://phylogatr.org/. An indicated shortcoming of current biodiversity data aggregators is the lack of back and forth communication between primary producers of data, data aggregators, and end-users (Anderson et al. 2020). We provide a means for submitting feedback and suggesting edits and data flags via a link to a Discourse page that is reviewed by the team of biologists and computer programmers.

## Empirical Example

We explored how genetic diversity is correlated with range size in almost 80,000 species and over 2 million sequences from the database (Table 1). Genetic diversity is an estimate of variation in the base pairs of DNA within a species, and is the basis for evolutionary change. Many measures of genetic diversity exist and can be used to understand different aspects about an individual, population, species, or community. By looking at patterns in genetic diversity, inferences can be made regarding evolutionary processes like migration, selection, and drift, and is often a first step in most genetic studies. Several measures of genetic diversity exist that capture different aspects of the data, such as estimates of the number of segregating sites (S), the number of haplotypes (H), and the mean per-site pairwise number of nucleotide differences sequences (π). It is expected that widespread species would have higher genetic diversity due to lower genetic drift and stable meta-population structure. Custom R v4.0.4 (R Core Team 2020) scripts were used to analyze data from several taxonomic groups by downloading sequence alignments by taxonomic group from the *phylogatR* database between 18-May-2021 and 11-June-2021.

**Table 1.**
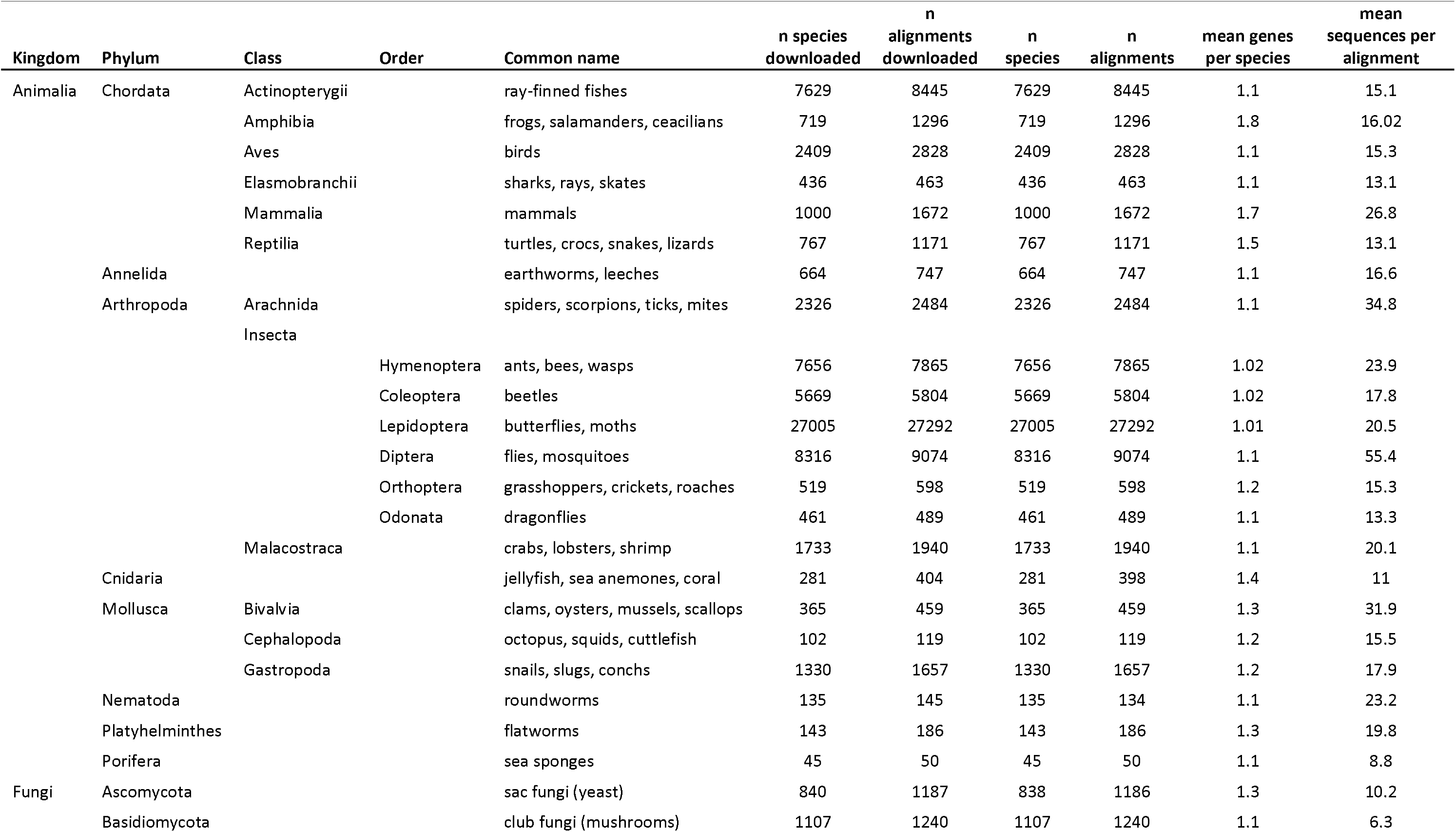
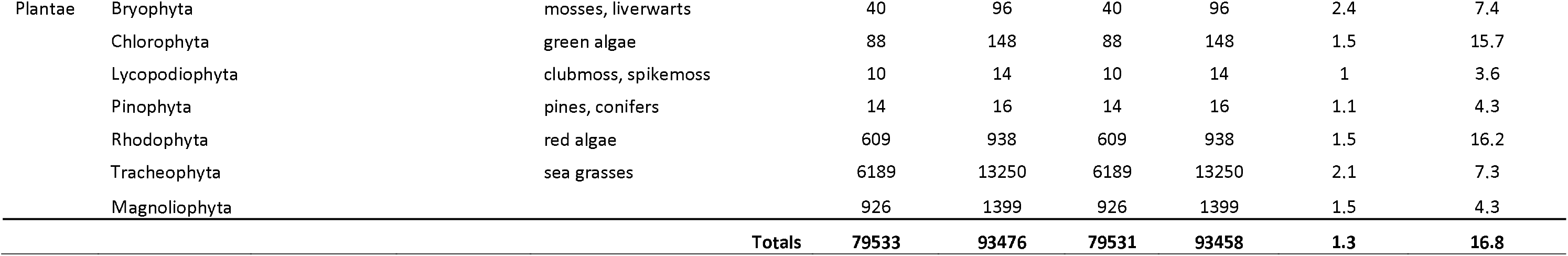
Summary of data downloaded from the database for analysis. Downloads were conducted by the lowest taxonomic group listed in the table. The number of species and alignments are those that were included in the data analysis pipeline before and after checking for binomial nomenclature and genetic or geographic outliers.

First, species names were scanned using the *genes.txt* files to find typos in species names, as well as other abnormalities in naming patterns. In several groups there were some non-binomial naming patterns (Supplemental Table S2). In the Platyhelminthes, Nematodes, Bivalves, Elasmobranchs, Hymenoptera, Lepidoptera, Diptera, Malacostraca, and Chlorophyta, there were several Genera that included species names labeled as letters and/or roman numerals (e.g., Co*tylurus c* or *f*.; *Paratylenchus BITH I* or *II*; *Hiatella C* or *D; Squalus clade B* or *C*; *Braunsapis A* or *B; Adoxophyes C* or *D*; *Allograpta CR A* or *B*; *Uristes murrayi morphospecies A* or *B; Ostreonium TeA* or *TeF*). In these instances, taxonomic expertise will be needed in deciding whether to treat these as different species. In one case there seemed to be an indication of a lateral gene transfer in Tracheophyta, which would need to be treated with caution (*Alloteropsis semialata* PCK 1P1 LGT:C and PPC 1P3 LGT:M). In another case, there was a misspelling in the name that we have updated in the database. This is an area of work where we are seeking user input but overall, the level of errors detected based on our exploration of these data are quite low, and easily checked by eye. The regression analysis below was carried out on the data with and without these abnormalities removed, and none had a significant impact on the results.

Nucleotide diversity (π) was calculated for each sequence alignment using the nuc.div function from the R package pegas (Paradis 2010). Geographic coordinates from each species were used to estimate the range of the species, though this only represents the sampling range of a species. Scatter plots of area and π were created for each taxonomic group using the package ggplot2 (Wickham 2017) to examine the data for outliers (Supplemental Table S3 and Supplemental Figures Zip File). When outliers were detected by area, online distribution maps were compared to the geographic coordinates from the dataset. In all these cases (58 total), the coordinates fell within the known published distributions. When outliers were detected by π (23 total), the geographic coordinates were also checked according to the published distributions. Again, no points fell outside the published distributions. These sequence alignments were also checked for possible mis-identified sequences or poorly aligned sequences. In most cases, a sequence or two slipped through our data cleaning steps and likely does not belong to either that species or locus and therefor produced a poor sequence alignment. The regression analysis below was conducted with and without the π and area outliers removed, and none had a significant impact on the results (Supplemental Table S3).

Several other sequence alignments from our initial download were not included in the following analyses (Supplemental Table S4). These alignments produced NA values for π (1050 total) and were explored further. In the majority (~95%) of cases, different portions of a given gene were sequenced such that there was no overlap in the middle of the sequence. In these instances, it is incumbent on the user to determine whether this level of missing data is appropriate for their analysis. The remaining cases were attributed to poor sequence alignments, usually due to just one sequence passing through our data cleaning steps. As such cases are discovered, alignments will be manually curated and updated in the database. As bad alignments are discovered, user input via the help documentation is encouraged.

Regression analysis was conducted to determine whether the size of geographic sampling could explain variation in genetic diversity using the lm function in R. Since we conducted 31 regression analyses, a Bonferroni correction was used to adjust our p-value (0.05 / 31 = 0.0016). Ten out of the 31 tests were significant (Table 2). In the vertebrates, only Actinopterygii, Elasmobranchii, and Mammalia were significant. In the insects, the Hymenoptera, Coleoptera, Lepidoptera, and Orthoptera, were significant. Porifera was significant, along with two plant groups (Rhodophyta and Tracheophyta). Porifera stands out as having a particularly high R-square value. Otherwise, no patterns emerge as far as which taxonomic groups would be more likely to display a relationship between area and π, or whether being winged, terrestrial, etc., for example, would contribute to an increase or decrease in genetic diversity, given the size of a species geographic distribution. There are likely a combination of factors that contribute to levels of genetic diversity within a species. This analysis is only a first step towards understanding how life history and dispersal ability may contribute to genetic variation and population structure.

**Table 2.** Summary of linear regression results. A Bonferroni correction was used to adjust our p-value (0.05 / 31 = 0.0016).

| **Group** | **pi**<br>**(mean)** | **R-**<br>**square** | **p-value** |
| --- | --- | --- | --- |
| Actinopterygii | 0.0165 | 0.0085 | **2.20E-16** |
| Amphibia | 0.0258 | -0.0007 | 0.776 |
| Aves | 0.0056 | -0.0014 | 0.8495 |
| Elasmobranchii | 0.0100 | 0.0274 | **0.0002** |
| Mammalia | 0.0218 | 0.0134 | **1.44E-06** |
| Reptilia | 0.0491 | -0.0002 | 0.3644 |
| Annelida | 0.0361 | 0.0041 | 0.0448 |
| Arachnida | 0.0150 | 0.0021 | 0.0034 |
| Hymenoptera | 0.0187 | 0.0020 | **5.09E-05** |
| Coleoptera | 0.0132 | 0.0038 | **5.74E-07** |
| Lepidoptera | 0.0104 | 0.0033 | **2.20E-16** |
| Diptera | 0.0140 | 0.0000 | 0.4218 |
| Orthoptera | 0.0147 | 0.0153 | **0.0014** |
| Odonata | 0.0398 | -0.0019 | 0.7434 |
| Malacostraca | 0.0279 | 0.0008 | 0.1069 |
| Cnidaria | 0.0376 | 0.0035 | 0.2394 |
| Bivalvia | 0.0245 | 0.0173 | 0.0027 |
| Cephalopoda | 0.0138 | 0.0208 | 0.0637 |
| Gastropoda | 0.0260 | -0.0006 | 0.8771 |
| Nematoda | 0.0219 | 0.0032 | 0.2276 |
| Platyhelminthes | 0.0270 | -0.0055 | 0.9588 |
| Porifera | 0.0141 | 0.7833 | **2.20E-16** |
| Ascomycota | 0.0132 | -0.0010 | 0.6397 |
| Basidiomycota | 0.0183 | 0.0054 | 0.0065 |
| Bryophyta | 0.0104 | -0.0092 | 0.7178 |
| Chlorophyta | 0.0349 | -0.0012 | 0.3605 |
| Lycopodiophyta | 0.1067 | -0.0504 | 0.4725 |
| Pinophyta | 0.1989 | 0.1868 | 0.0534 |
| Rhodophyta | 0.0081 | 0.0258 | **5.24E-07** |
| Tracheophyta | 0.0236 | 0.0109 | **2.20E-16** |
| Magnoliophyta | 0.0394 | -0.0004 | 0.5011 |

Two plant groups have relatively high values for π (e.g., Lycopodiophyta, Pinophyta). This suggests these groups are worth further exploration, as either they may be in need of database updates to reflect taxonomic revisions and misidentifications, or these groups may harbor a high number of cryptic species (Parsons et al. In review). Documenting global levels of genetic diversity, an important measure of biodiversity, can serve as a baseline for detecting rapid changes, or loss of diversity, due to climate change (Paz-Vinas et al. 2018). Furthermore, measures of genetic variation are often used to assess a species’ or population’s ability to respond to environmental (climate, habitat, biotic) changes (Frankham 2005; Hoffman and Sgró 2011), so large-scale analyses such as this, allow for targeting individual species that might be at a higher risk for extinction (Frankham et al. 2014; Hoban et al. 2020) and for identifying species attributes that contribute to higher levels of genetic diversity (Broadhurst et al. 2017). While there is no consensus as to whether measures of genetic diversity from a single mitochondrial or chloroplast gene, the most common in our dataset, are appropriate measures of genetic diversity (Petit-Marty et al. 2021; Paz-Vinas et al. 2021), many species (15%; 13,960 total) have data from multiple loci in *phylogatR* and measures of genetic diversity across loci can be evaluated. However, including spatial information for individuals allows further insight into the factors that contribute to increasing or decreasing levels of genetic variation, such as shared barriers to dispersal and responses to environmental change. Genetic diversity alone may not be a strong indicator of species stability but integrating the information that can be gained via geographic coordinates (e.g., climate layers) is necessary to consider demographic history and environmental variables for implementing effective conservation strategies (Teixeira & Huber 2021).

A useful secondary product of the analysis described above is the opportunity to explore outliers and inconsistencies in the database. We identified 1131 alignments (1.2% of the data) that could potentially bias our results. While in our case there is sufficient data that a small amount of noise caused by outliers and inconsistent species names did not influence the results (Supplemental Tables S2-S4), this may not be universally true for all analyses. We had 1,511,882 occurrences with flags (Table S5). Of those that were flagged, the majority of these were flag ‘**g**’ (50%), followed by flag ‘**d**’ (18%), suggesting many historical DNA sequences had been uploaded to GenBank as haplotypes. We recommend those uploading data to these databases refrain from uploading haplotypic data and include DNA sequences from all individuals. Flag ‘**b**’ (26%) and flag ‘**m**’ (4.8%) were the next most common flags, suggesting there are many duplicates in these databases that need to be removed. Finally, flag ‘**s**’ occurred in only 0.2% of flagged occurrences, indicating that taxonomy issues are present, but do not overwhelm the data. Users of *phylgatR* are cautioned to pay attention to flagged sequences and alignments and to make appropriate corrections as dictated by the needs of their investigation protocol. The scripts used for these analyses are available on the *phylogatR* website and can be used to facilitate screening the data.

The exploration of these data began in a bioinformatics course that aimed to introduce students to multiple sequence alignments, highlight the value of estimating genetic diversity and using open-source databases, and learn the structure of creating loops. This work resulted in an undergraduate researcher from the course leading the efforts for this empirical example and being co-author on this paper. The datasets that can be generated via *phylogatR* will contribute to the ongoing development of resources that will expose students to real data and computational methods in the classroom. Incorporating authentic research into classroom instruction provides inclusive learning experiences for all students and leads to better learning outcomes (Theobald et al. 2020). The additional benefit of *phylogatR* is that concepts in evolution and ecology can be taught with real data at no cost, other than computer access. The *phylogatR* website contains teacher resources, which include teaching modules and associated instructor notes, with intent to increase these resources in the future.

## Conclusions

Identifying the evolutionary and environmental processes that have influenced a single lineage is an ongoing challenge for evolutionary biologists, but a true understanding of these processes will require the synthesis of results from thousands of individual studies. Such a synthesis will be most efficiently achieved via data repurposing and automated analysis. *phylogatR* makes such syntheses more accessible for all researchers. By bypassing the idiosyncratic results of individual studies, *phylogatR* will enable biologists to test hypotheses at various taxonomic and geographic scales. The example analysis presented above combines genetic and geographic data in a way that is only meaningful when done on a large scale. These results indicate that we will make fundamental contributions to understanding global patterns of genetic diversity that will have important implications to conservation management and species discovery (see Table 3).

**Table 3.** The *phylogatR* database and web portal can enable the testing of these hypotheses (among others) on a continental or global scale.

| Hypothesis | Example Reference |
| --- | --- |
| Current ecological communities are historically stable | Zink 2002 |
| Shared organismal traits lead to concordant phylogeographic patterns | Papadopoulou & Knowles 2015; Zamudio et al. 2016 |
| Members of ecologically-interdependent communities will co-diversify | Smith et al. 2011; Satler & Carstens 2016 |
| Pleistocene refugia are shared by species from many taxonomic groups | Brunsfeld et al. 2001 |
| Cosmopolitan species will have higher levels of genetic diversity than small endemics | Gitzendanner & Soltis 2000 |
| Regions of marginal habitat contain less genetic diversity | Micheletti & Storfer 2015 |
| Generalist species will have weaker responses to climatic and landscape changes than habitat specialists | Estavillo et al. 2013 |
| Southern peninsular regions served as Pleistocene refugia in the Northern Hemisphere | Hewitt 1996 |
| Cryptic species are likely to be present in regions of high endemism | Reeder et al. 2007 |
| Ecological niche differentiation promotes genetic diversification | McCormack et al. 2010 |

While single-locus data has its limitations in making inferences about historical demography (Matumba et al. 2020), DNA barcoding, or the use of other single-locus DNA markers has provided tremendous insight into identifying evolutionary significant units and providing information on species in further need of exploration (Sholihah et al. 2020; Wang et al. 2020; Nneji et al. 2020; Bousjein et al. 2021; León-Tapia 2021). These data are particularly helpful when aiming to explore broad-scale patterns such as those on a continental scale (Dinça et al. 2021) or across species (Doorenweerd et al. 2020), especially for a large number of taxonomic groups, as demonstrated here. Studies using data from our initial data aggregation pipelines have further demonstrated the utility of single-locus large-scale studies. Parsons et al. (In review) explore cryptic diversity in mammals using molecular species delimitation methods for single-locus data in conjunction with natural history and environmental data for over 4,000 species. They found that hundreds of mammal species are still likely undescribed and that these are mostly small-bodied taxa with large ranges. Fonesca et al. (In review) examined intraspecific genetic diversity in the context of the climate stability hypothesis for over 30,000 species. This work suggests that population bottlenecks and range expansions in the temperate regions around the world have lasting effects on genetic diversity, resulting in significantly higher intraspecific genetic diversity in the tropics.

Our brief empirical example allowed us to document outliers in the data and search for poor sequence alignments, as we will continue to improve the database and data curation steps. We will continue to make recommendations and supply users with guidance in data checking before analysis. We encourage continued natural history work to better populate biodiversity databases as the benefits of publicly available data are numerous and experts are needed to correct database errors and decide where data deficiencies lie (Groom et al. 2020; Leigh et al. 2021). Further, making data easy to access and reuse is important for researchers and educators who do not have the skills or resources for large-scale projects, or expensive and time-consuming field and lab work, increasing participation from underprivileged groups and minorities (Estrada et al. 2016; Whittington & Pelletier 2021). By making real genetic data available to students from any school with a connection to the internet, *phylogatR* will inspire the next generation of researchers to understand and protect biodiversity while they are developing the computational skills that are increasingly required for evolutionary and ecological studies. Not only do these data make authentic research more readily available in the classroom, they increase the access to biodiversity data worldwide, therefore contributing to a more inclusive and diverse STEM community and easily implemented international collaborations (Heberling et al. 2021; Marden et al. 2021).

Perfect data is unattainable and not all data will be retained after data curations steps (Peterson et al. 2018). The data currently available on *phylogatR* offer a first step towards asking big questions with big data in population genetics, phylogeography, and systematics. While this study does not aim to solve problems in data standards, making data more readily available will likely result in novel questions and transformative findings, and will contribute to identifying current shortcomings and inconsistencies in current data sharing practices.

## Acknowledgements

We thank the several OSC members for their participation in database development, Alan Chalker, and the *phylogatR* beta-testers group for assessing functionality of the database. Funding was provided by the National Science Foundation (DBI-1910623) to BCC and the National Science Foundation (DBI-1911293) to TAP.

## Data Accessibility and Benefit-Sharing Statement

The phylogatR database is publicly available at https://phylogatr.org/ where every download includes the GBIF DOI, GenBank version, and BOLD DOI that contributed to the data. All scripts devoted to the development of the database can be found at https://github.com/OSC/phylogatr-web. Benefits Generated: Benefits from this research include accessibility to big data via the public database, minimizing the need for computational resources, as described above, which include data analysis pipelines and educational tools.

## Author Contributions

TAP: designed research, performed research, wrote paper, acquired funding. BCC: designed research, performed research, wrote paper, acquired funding. DP: performed research, edited paper, created figures. SD: performed research, edited paper, created logo. SC: analyzed data, edited paper. EF: contributed code. JO: contributed code.

## References

Anderson, R. P., Araújo, M. B., Guisan, A., Lobo, J. M., Martínez-Meyer, E., Peterson, A. T., & Soberón, J. M. (2020). Optimizing biodiversity informatics to improve information flow, data quality, and utility for science and society. Frontiers of Biogeography, 12(3), e47839. https://doi.org/10.21425/F5FBG47839

Blanchet, S., Prunier, J. G., & De Kort, H. (2017). Time to Go Bigger: Emerging Patterns in Macrogenetics. Trends in Genetics, 33(9), 579–580. https://doi.org/10.1016/j.tig.2017.06.007

Bousjein, N. S., Gardner, M. G., & Schwarz, M. P. (2021) Demographic stability of the Australian temperate exoneurine bees (Hymenoptera: Apidae) through the Last Glacial Maximum. Austral Entomology, 60(3), 549–559. https://doi.org/10.1111/aen.12539

Broadhurst, L., Breed, M., Lowe, A., Bragg, J., Catullo, R., Coates, D., … Byrne, M. (2017). Genetic diversity and structure of the Australian flora. Diversity and Distributions, 23(1), 41–52. https://doi.org/10.1111/ddi.12505

Brunsfeld, S. J., Sullivan, J., Soltis, D. E., & Soltis, P. S. (2000). Comparative phylogeography of north-western North America: a synthesis. In: Silvertown J, Antonovics J, editors. Integrating ecology and evolution in a spatial context. Williston, VT: Blackwell Publishing, p 319–339.

Burbrink, F. T., Gehara, M., McKelvy, A. D., & Myers, E. A. (2021). Resolving spatial complexities of hybridization in the context of the gray zone of speciation in North American ratsnakes (*Pantherophis obsoletus* complex). Evolution, 75(2), 260 277. https://doi.org/10.1111/evo.14141

Carstens, B. C., Morales, A. E., Field, K., & Pelletier, T. A. (2018). A global analysis of bats using automated comparative phylogeography uncovers a surprising impact of Pleistocene glaciation. Journal of Biogeography, 45(8), 1795–1805. https://doi.org/10.1111/jbi.13382

Cavalcante, T., Jesus, A. S., Rabelo, R. M., Messias, M. R., Valsecchi, J., Ferraz, … Barnett, A. A. (2020). Niche overlap between two sympatric frugivorous Neotropical primates: Improving ecological niche models using closely-related taxa. Biodiversity and Conservation, 29(8), 2749–2763. https://doi.org/10.1007/s10531-020-01997-5

Chamberlain, S. A., & Szöcs, E. (2013). taxize: taxonomic search and retrieval in R. F1000Research, 2, 191. https://doi.org/10.12688/f1000research.2-191.v2

Dawson, M. N. (2014). Natural experiments and meta-analyses in comparative phylogeography. Journal of Biogeography, 41(1), 52 65. https://doi.org/10.1111/jbi.12190

Dincă, V., Dapporto, L., Somervuo, P., Vodă, R., Sylvain Cuvelier, Martin Gascoigne-Pees, Huemer, P., Mutanen, M., Hebert, P. D. N., & Vila, R. (2021). High resolution DNA barcode library for European butterflies reveals continental patterns of mitochondrial genetic diversity. Communications Biology, 4(315). https://doi.org/10.1038/s42003-021-01834-7

Doorenweerd, C., San Jose, M., Barr, N., Leblanc, L., & Rubinoff, D. (2020). Highly variable COI haplotype diversity between three species of invasive pest fruit fly reflects remarkably incongruent demographic histories. Scientific Reports, 10(1), 6887. https://doi.org/10.1038/s41598-020-63973-x

Estavillo, C., Pardini, R., & da Rocha, P. L. B. (2013). Forest loss and the biodiversity threshold: An evaluation considering species habitat requirements and the use of matrix habitats. Plos One, 8(12), e82369. https://doi.org/10.1371/journal.pone.0082369

Estrada, M., Burnett, M., Campbell, A. G., Campbell, P. B., Denetclaw, W. F., Gutiérrez, C. G., … Zavala, M. (2016). Improving Underrepresented Minority Student Persistence in STEM. CBE life sciences education, 15(3), es5. https://doi.org/10.1187/cbe.16-01-0038

Farallo, V. R., Muñoz, M. M., Uyeda, J. C., & Miles, D. B. (2020). Scaling between macro- to microscale climatic data reveals strong phylogenetic inertia in niche evolution in plethodontid salamanders. Evolution, 74(5), 979–991. https://doi.org/10.1111/evo.13959

Folk, R. A., & Siniscalchi C. M. (2021). Biodiversity at the global scale: The synthesis continues. American Journal of Botany, 108(6), 912–924. https://bsapubs.onlinelibrary.wiley.com/doi/full/10.1002/ajb2.1694

Fonesca, E. M., Pelletier, T. A., Decker, S. K., Parsons, D. J., Carstens, B. C. Quaternary climate oscillations caused the latitudinal gradient of intraspecific genetic diversity. In Review.

Frankham, R. (2005). Genetics and extinction. Biological Conservation, 126(2), 131–140. https://doi.org/10.1016/j.biocon.2005.05.002

Frankham, R., Bradshaw, C. J. A., & Brook, B. W. (2014). Genetics in conservation management: Revised recommendations for the 50/500 rules, Red List criteria and population viability analyses. Biological Conservation, 170, 56–63. https://doi.org/10.1016/j.biocon.2013.12.036

Garrick, R. C., Bonatelli, I. A. S., Hyseni, C., Morales, A., Pelletier, T. A., Perez, M. F., … Carstens, B. C. (2015). The evolution of phylogeographic data sets. Molecular Ecology, 24(6), 1164–1171. https://doi.org/10.1111/mec.13108

Gratton, P., Marta, S., Bocksberger, G., Winter, M., Trucchi, E., & Kühl, H. (2017). A world of sequences: Can we use georeferenced nucleotide databases for a robust automated phylogeography?. Journal of Biogeography, 44(2), 475–4486. https://doi.org/10.1111/jbi.12786

Gitzendanner, M. A., & Soltis, P. S. (2000). Patterns of genetic variation in rare and widespread plant congeners. American Journal of Botany, 87(6), 783–792. https://doi.org/10.2307/2656886

Groom, G., Güntsch, A., Huybrechts, P., Kearney, N., Leachman, S., Nicolson, N., Page, R. D. M., Shorthouse, D. P., Thessen, A. E., & Haston, E. (2020). People are essential to linking biodiversity data. Database, 2020, baaa072. https://doi.org/10.1093/database/baaa072

Guralnick, R., & Hill, A. (2009). Biodiversity informatics: automated approaches for documenting global biodiversity patterns and processes. Bioinformatics, 25(4), 421–428. https://doi.org/10.1093/bioinformatics/btn659

Hickerson, M. J., Carstens, B. C., Cavender-Barnes, K. A., Crandall, C. H., Graham, J. B., Johnson, L., … Yoder, A. D. (2010). Phylogeography’s past, present, and future: 10 years after Avise, 2000. Molecular Phylogenetics and Evolution, 54(1), 291–301. https://doi.org/10.1016/j.ympev.2009.09.016

Hewitt, G. M. (1996). Some genetic consequences of ice ages, and their role in divergence and speciation, Biological Journal of the Linnean Society, 58(3), 247–276. https://doi.org/10.1111/j.1095-8312.1996.tb01434.x

Heberling, J. M., Miller, J. T., Noesgaard, D., Weingart, S. B., & Schigel, D. (2021). Data integration enables global biodiversity synthesis. Proceedings of the National Academy of Sciences, 118(6). https://www.pnas.org/content/118/6/e2018093118/tab-article-info

Hoban, S., Bruford, M., Jackson, J. D’U, Lopes-Fernandes, M., Heuertz, M., Hohenlohe, P. A., … Laikre, L. (2020). Genetic diversity targets and indicators in the CBD post-2020 Global Biodiversity Framework must be improved. Biological Conservation, 248, 108654. https://doi.org/10.1016/j.biocon.2020.108654

Hoffmann, A., & Sgrò, C. (2011). Climate change and evolutionary adaptation. Nature, 470, 479–485. https://doi.org/10.1038/nature09670

Leigh, D. M., van Rees, C. B., Millette, K. L., Breed, M. F., Schmidt, C., Bertola, L. D., … Paz-Vinas, I. (2021). Opportunities and challenges of macrogenetic studies. Nature Reviews Genetics. https://doi.org/10.1038/s41576-021-00394-0

León-Tapia, M. Á. (2021). DNA barcoding and demographic history of *Peromyscus yucatanicus* (Rodentia: Cricetidae) endemic to the Yucatan peninsula, Mexico. Journal of Mammalian Evolution, 28(2), 481–495. https://doi.org/10.1007/s10914-020-09510-z

Marden, E., Abbott, R. J., Austerlitz, F., Ortiz-Barrientos, D., Baucom, R. S., Bongaerts, P., … Rieseberg, L. H. (2021). Sharing and reporting benefits from biodiversity research. Molecular Ecology, 30(5), 1103–1107. https://doi.org/10.1111/mec.15702

Marques AC, Maronna, M. M., Collins, A. G. (2013). Putting GenBank data on the map. Science, 341(6152), 1341. https://www.science.org/doi/10.1126/science.341.6152.1341-a

Matumba, T. G., Oliver, J, Barker, N. P., McQuaid, C. D., & Teske, P. R. (2020). Intraspecific mitochondrial gene variation can be as low as that of nuclear rRNA. F1000Research, 9, 339. https://doi.org/10.12688/f1000research.23635.2

Micheletti, S. J., & Storfer, A. (2015). A test of the central-marginal hypothesis using population genetics and ecological niche modelling in an endemic salamander (*Abystoma barbouri)*. Molecular Ecology, 24(5), 967–979. https://doi.org/10.1111/mec.13083

Miraldo, A., Li, S., Borregaard, M. K., Flórez-Rodiríguez, A., Gopalakrishnan, S., Rizvanovic, M., Wang, Z., … Nogués-Bravo, D. (2016). An Anthropocene map of genetic diversity. Science, 353(6307), 1532–1535. https://pubmed.ncbi.nlm.nih.gov/27708102/

Nneji, L., M., Adeola, A. C., Mustapha, M. K., Oladipo, S. O., Djagoun, C. A. M. S., Nneji, I. C., … Nwani, C. D. (2020). DNA barcoding silver butterfish (*Schilbe intermedius*) reveals patterns of mitochondrial genetic diversity across African river systems. Scientific Reports, 10(7097). https://doi.org/10.1038/s41598-020-63837-4

Nottingham, S., Pelletier, T. A. (2021). The impact of climate change on western *Plethodon* salamanders’ distribution. Ecology and Evolution, 11(14), 9370–9384. https://doi.org/10.1002/ece3.7735

Papadopoulou, A., & Knowles, L. L. (2015). Species-specific responses to island connectivity cycles: Refined models for testing phylogeographic concordance across a Mediterranean Pleistocene aggregate island complex. Molecular Ecology, 24(16), 4252–4268. https://doi.org/10.1111/mec.13305

Papadopoulou, A., & Knowles, L. L. (2016). Toward a paradigm shift in comparative phylogeography driven by trait-based hypotheses. Proceedings of the National Academy of Sciences, 113(29), 8018–8024. https://doi.org/10.1073/pnas.1601069113

Paradis, E. (2010). pegas: an R package for population genetics with an integrated-modular approach. Bioinformatics, 26(3), 419–420. https://academic.oup.com/bioinformatics/article/26/3/419/215731

Parsons, D., Pelletier, T. A., Duckett, D., Wieringa, J., & Carstens, B. C. Analysis of biodiversity data suggest that species are hidden in predictable places. In review.

Paz-Vinas, I., Loot, G., Hermoso, V., Veyssière, C., Poulet, N., Grenouillet, G., & Blanchet, S. (2018). Systematic conservation planning for intraspecific genetic diversity. Proceedings of the Royal Society B: Biological Sciences, 285(1877), 2746. https://royalsocietypublishing.org/doi/10.1098/rspb.2017.2746

Paz-Vinas, I., Jensen, E. L., Bertola, L. D., Breed, M. F., Hand, B. K., Hunter, M. E., Kershaw, F., Leigh, D. M., Luikart, G., Mergeay, J., Miller, J. M., Van Rees, C. B., Segelbacher, G. & Hoban, S. (2021). Macrogenetic studies must not ignore limitations of genetic markers and scale. Ecology Letters, 24(6), 1282–1284. https://doi.org/10.1111/ele.13732

Pelletier, T. A., & Carstens, B. C. (2016). Comparing range evolution in two western *Plethodon* salamanders: glacial refugia, competition, ecological niches, and spatial sorting. Journal of Biogeography, 43(11), 2237–2249. https://doi.org/10.1111/jbi.12833

Pelletier, T. A., & Carstens, B. C. (2018). Geographical range size and latitude predict population genetic structure in a global survey. Biology Letters, 14(1), e20170566. https://royalsocietypublishing.org/doi/10.1098/rsbl.2017.0566

Pelletier, T. A., Carstens, B. C., Tank, D. C., Sullivan, J., & Espíndola, A. (2018). Predicting plant conservation priorities on a global scale. Proceedings of the National Academy of Sciences, 115(51), 13027–13032. https://www.pnas.org/content/115/51/13027

Peterson, A. T., Asase, A., Canhos, D., de Souza, S., & Wieczorek, J. (2018). Data Leakage and Loss in Biodiversity Informatics. Biodiversity data journal, 6, e26826. https://doi.org/10.3897/BDJ.6.e26826

Petit-Marty, N., Vázquez-Luis, M., & Hendriks, I. E. (2021). Use of the nucleotide diversity in COI mitochondrial gene as an early diagnostic of conservation status of animal species. Conservation Letters, 14, e12756. https://doi.org/10.1111/conl.12756

R Core Team. (2020). R version 4.0.2 – “Taking off Again”. The R Foundation for Statistical Computing.

Reeder, D. M., Helgen K. M., & Wilson, D. E. (2007). Global trends and biases in new mammal species discoveries. Occasional Papers Museum of Texas Tech University, 269, 1–36. https://www.biodiversitylibrary.org/item/263303#page/1/mode/1up

Satler, J. D., & Carstens, B. C. (2016). Phylogeographic concordance factors quantify phylogeographic congruence among co-distributed in the *Sarracenia alata* pitcher plant system. Evolution, 70(5), 1105–1119. https://doi.org/10.1111/evo.12924

Sholihah, A., Delrieu-Trottin, E., Sukmono, T., Dahruddin, H., Risdawati, R., Elvyra, R., … Hubert, N. (2020). Disentangling the taxonomy of the subfamily Rasborinae (Cypriniformes, Danionidae) in Sundaland using DNA barcodes. Scientific Reports, 10, 2818. https://doi.org/10.1038/s41598-020-59544-9

Sidlauskas, B., Ganapathy, G., Hazkani-Covo, E., Jenkins, K. P., Lapp, H., McCall, L. W., Price, S., Scherle, R., Spaeth, P. A. & Kidd, D. M. (2010). Linking big: The continuing promise of evolutionary synthesis. Evolution, 64(4), 871–880. https://doi.org/10.1111/j.1558-5646.2009.00892.x

Smith, C. I., Tank, S., Godsoe, W., Levenick, J., Strand, E., Esque, T., & Pellmyr, O. (2011). Comparative phylogeography of a coevolved community: Concerted population expansions in Joshua trees and four yucca moths. Plos One, 6(10), e25628. https://doi.org/10.1371/journal.pone.0025628

Smith, M. L. & Carstens, B. C. (2020). Process-based species delimitation leads to identification of more biologically relevant species. Evolution, 74(2), 216–229. https://doi.org/10.1111/evo.13878

Teixeira, J. C., & Huber, C. D. (2021). The inflated significance of neutral genetic diversity in conservation genetics. Proceedings of the National Academy of Sciences, 118(10). https://www.pnas.org/content/118/10/e2015096118

Theobald, E. J., Hill, M. J., Tran, E., Agrawal, S., Arroyo, E. N., Behling, S., … Scott Freeman. (2020). Active learning narrows achievement gaps for underrepresented students in undergraduate science, technology, engineering, and math. Proceedings of the National Academy of Sciences, 117(12), 6476–6483. https://www.pnas.org/content/117/12/6476

Thompson, C. E. P., Pelletier, T. A., & Carstens, B. C. (2021). Genetic diversity of North American vertebrates in protected areas. Biological Journal of the Linnean Society, 132(2), 388–399. https://doi.org/10.1093/biolinnean/blaa195

Wang, T., Zhang, Y-p., Yang, Z-y., Lui, Z., & Du, Y-y. (2020). DNA barcoding reveals cryptic diversity in the underestimated genus Triplophysa (Cypriniformes: Cobitidae, Nemacheilinae) from the northeastern Qinghai-Tibet Plateau. BMC Evolutionary Biology, 20, 151. https://doi.org/10.1186/s12862-020-01718-0

Whittington, A., & Pelletier, T. A. (2021). Women in Field Science: Challenges, Strategies and Supports for Success. Journal of Women and Minorities in Science and Engineering 27(6), 59–83. https://www.dl.begellhouse.com/journals/00551c876cc2f027,004d48e6514cf1a9,0fb2a5fa146133b3.html

Whitlock, M. C., McPeek, M. A., Rausher, M. D., Rieseberg, L., & Moore, A. J. (2010). Data Archiving. The American Naturalist, 175(2), 145–146. https://www.journals.uchicago.edu/doi/full/10.1086/650340

Wickham, E. (2017). ggplot2: Elegant Graphics for Data Analysis. Statistical Software, 77, b02. https://www.jstatsoft.org/v77/b02/

Wilkinson, M., Dumontier, M., Aalbersberg, I. Appleton, G., Axton, M., Baak, A., … Mons, B. (2016). The FAIR Guiding Principles for scientific data management and stewardship. Scientific Data, 3, 160018. https://doi.org/10.1038/sdata.2016.18

Zamudio, K. R, Bell, R. C., & Mason, N. A. (2016). Phenotypes in phylogeography: Species’ traits, Environmental variation, and vertebrate diversity. Proceedings of the National Academy of Sciences, 113(29), 8041–8048. https://www.pnas.org/content/113/29/8041

Zink, R., M. (2002). Methods in comparative phylogeography, and their application to studying evolution in the north American aridlands. Integrative and Comparative Biology, 42(5), 953–959. https://doi.org/10.1093/icb/42.5.953

