## Supplementary figures and images for "*phylogatR*: Phylogeographic data aggregation and repurposing"

### actinopterygii_plot.pdf

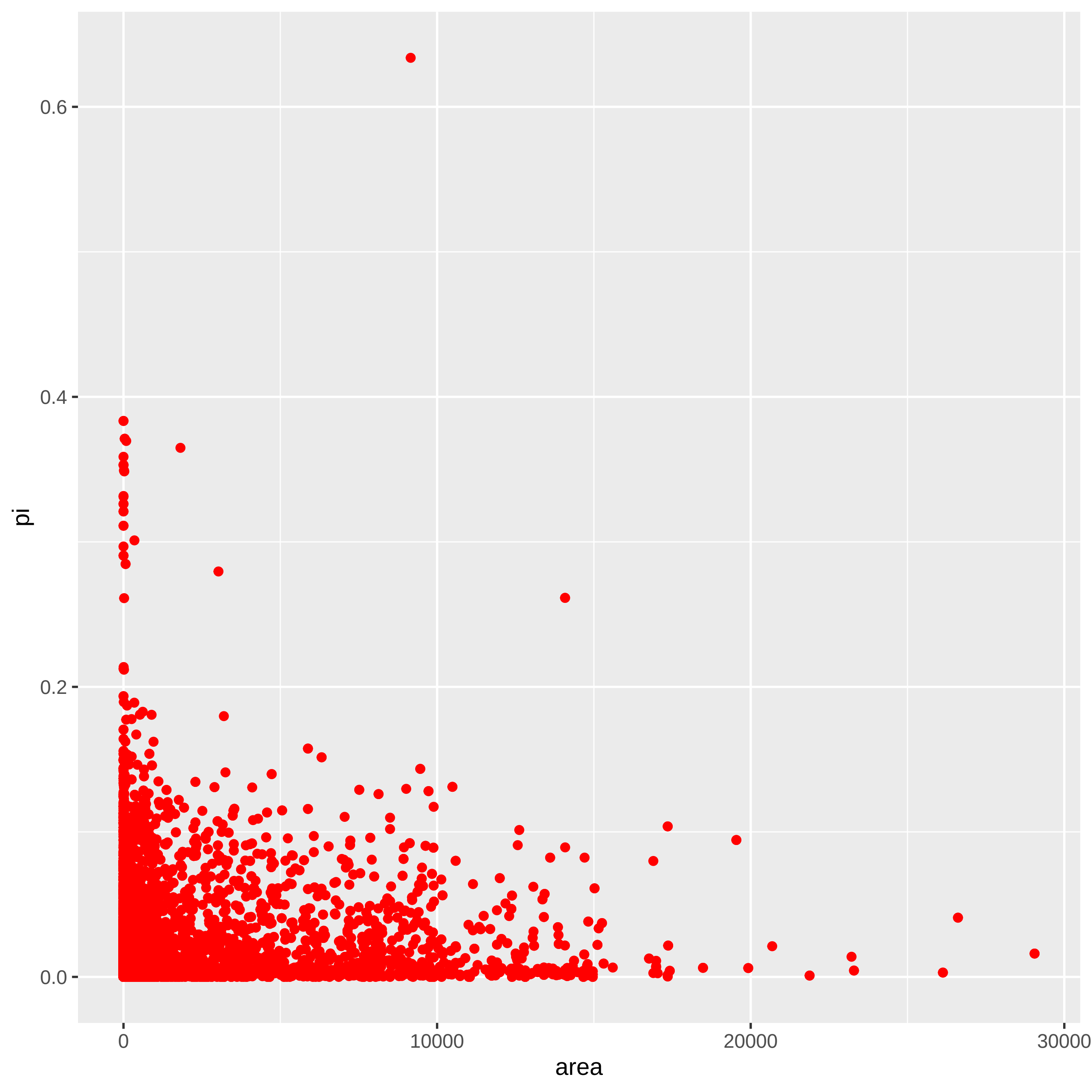

### amphibia_plot.pdf

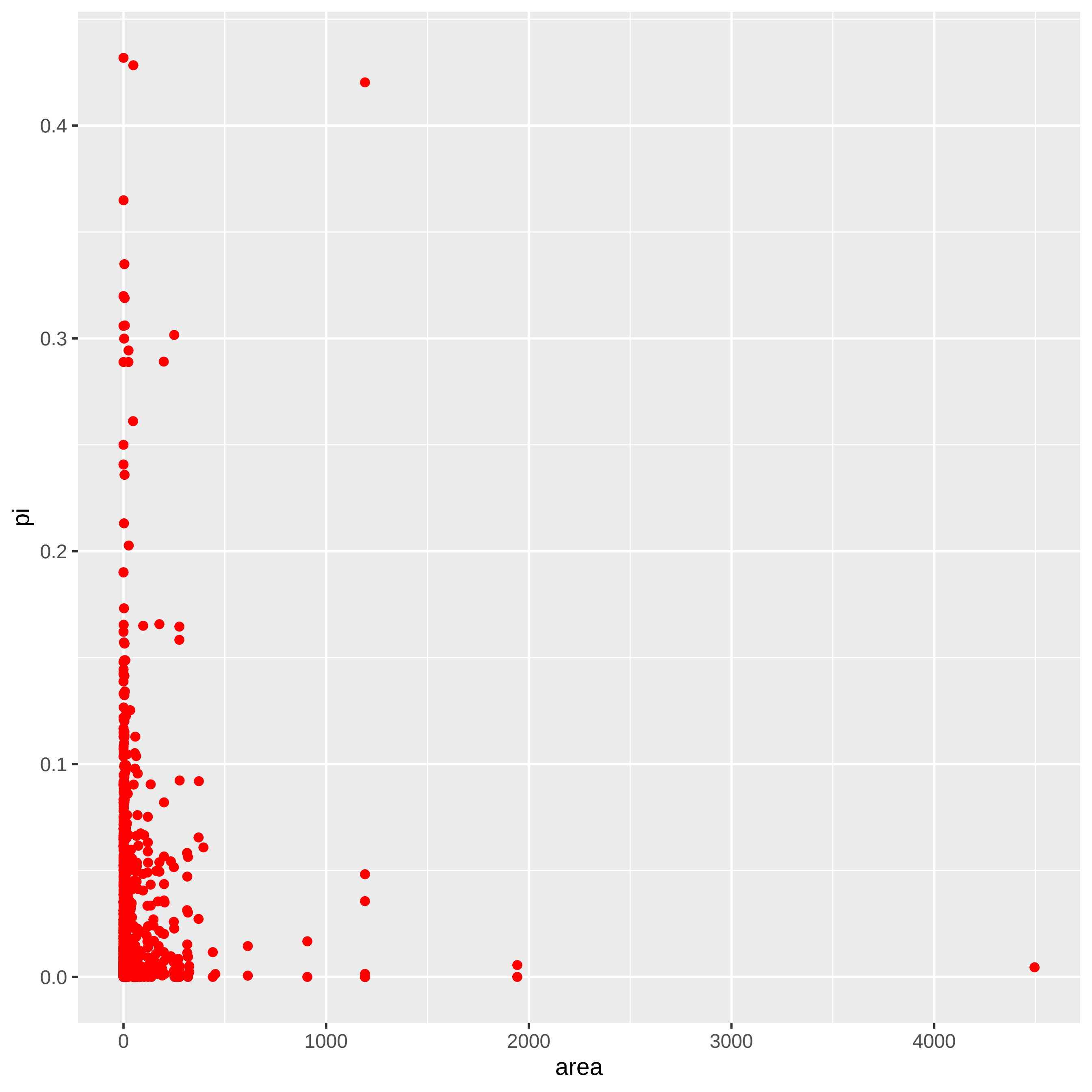

### annelida_plot.pdf

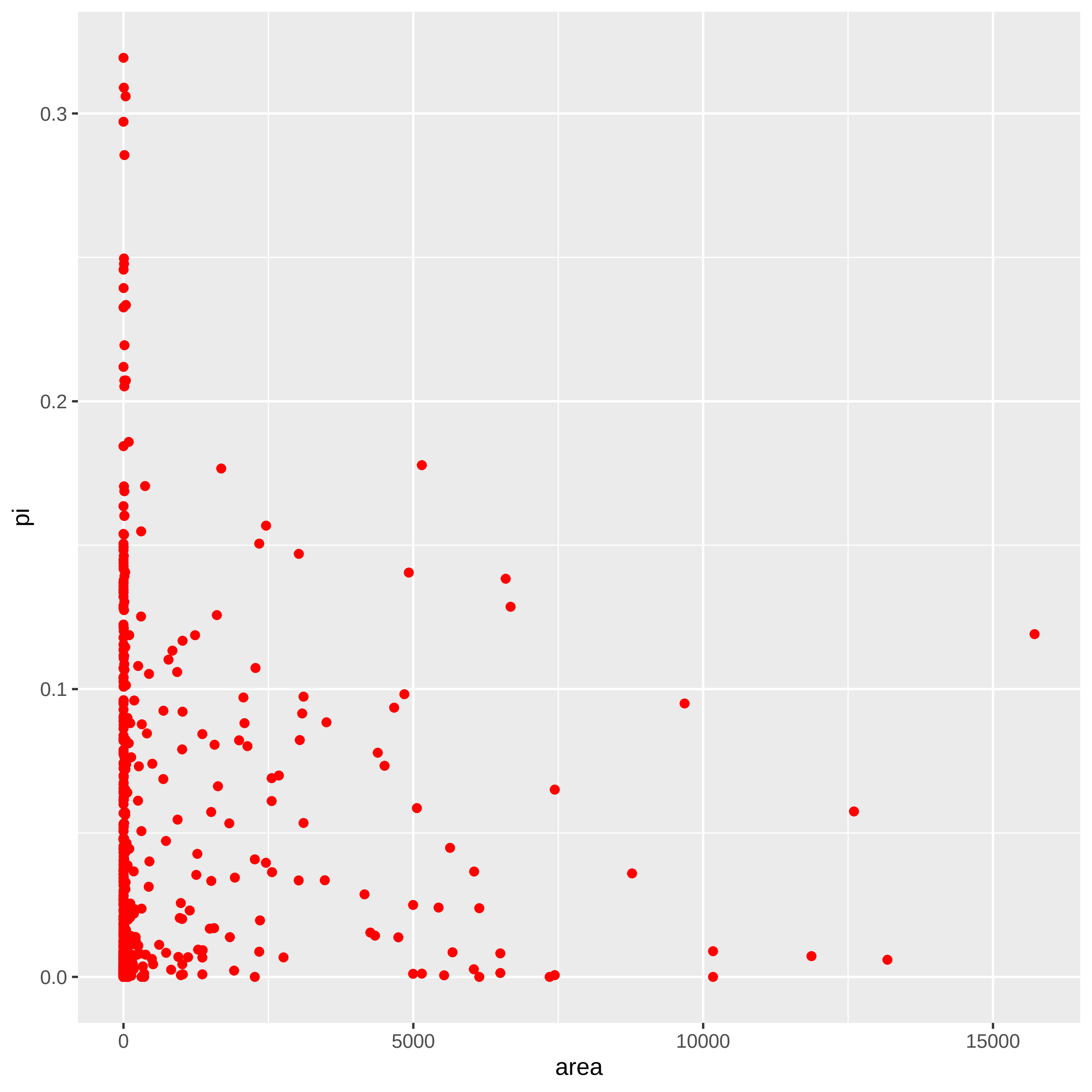

### arachnida_plot.pdf

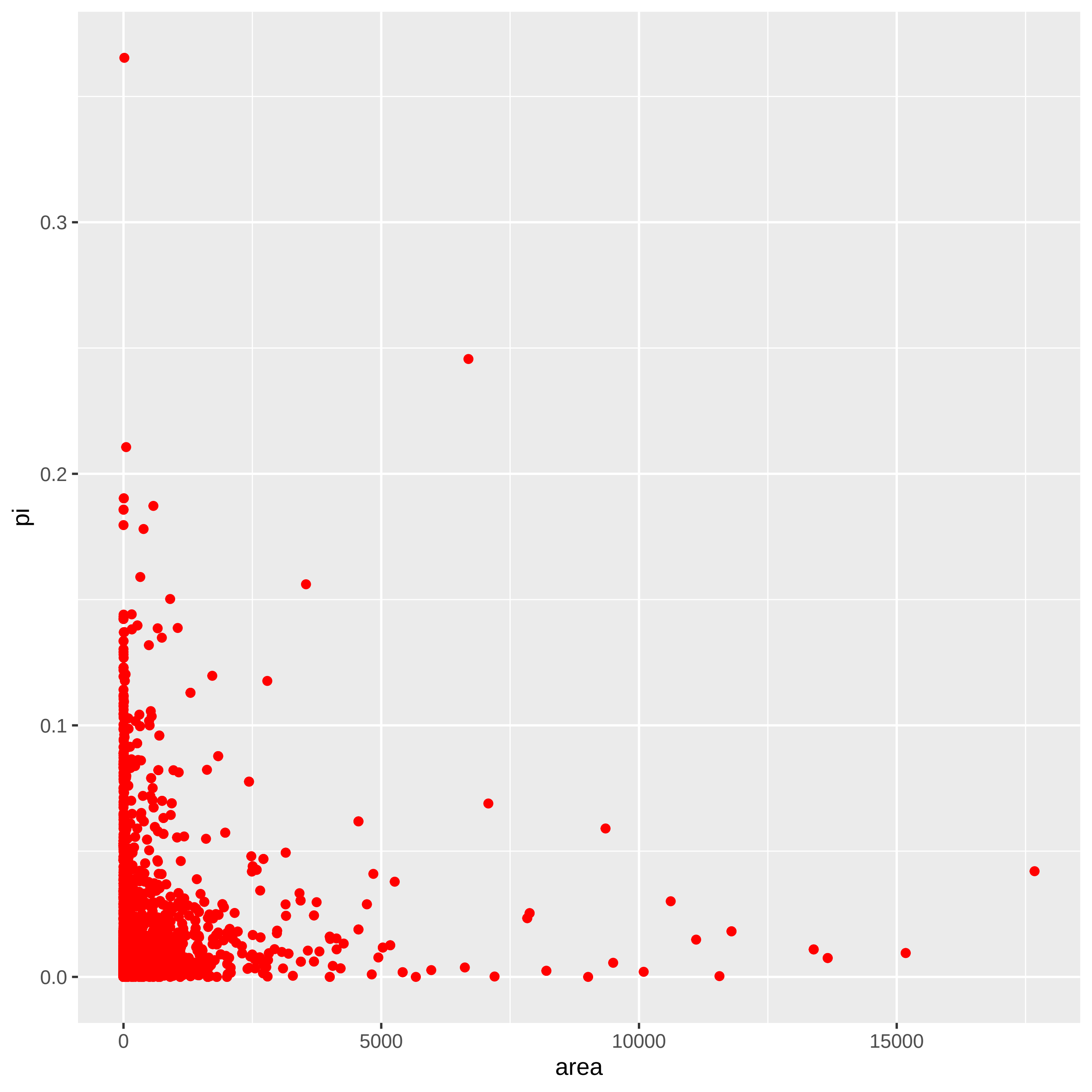

### ascomycota_plot.pdf

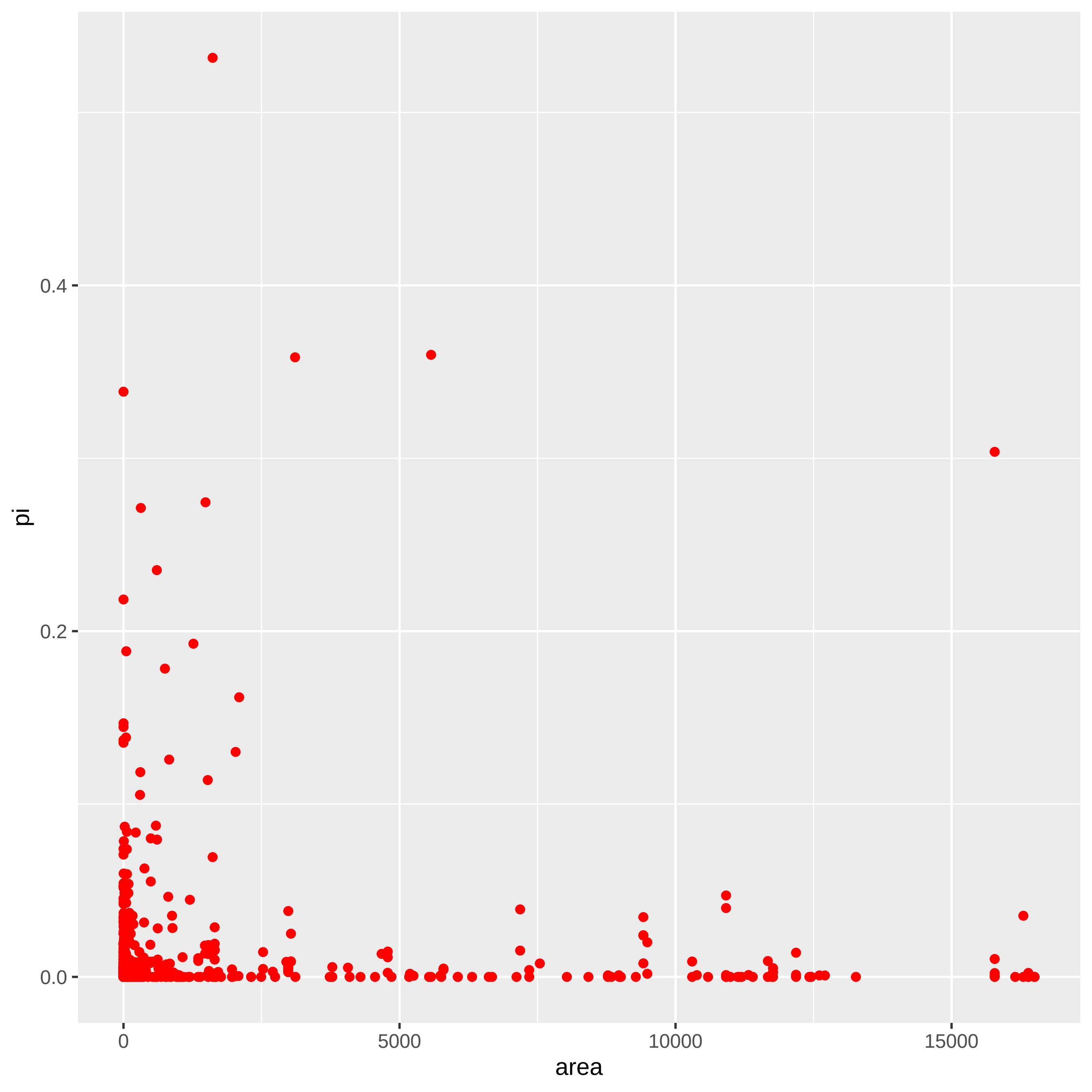

### aves_plot.pdf

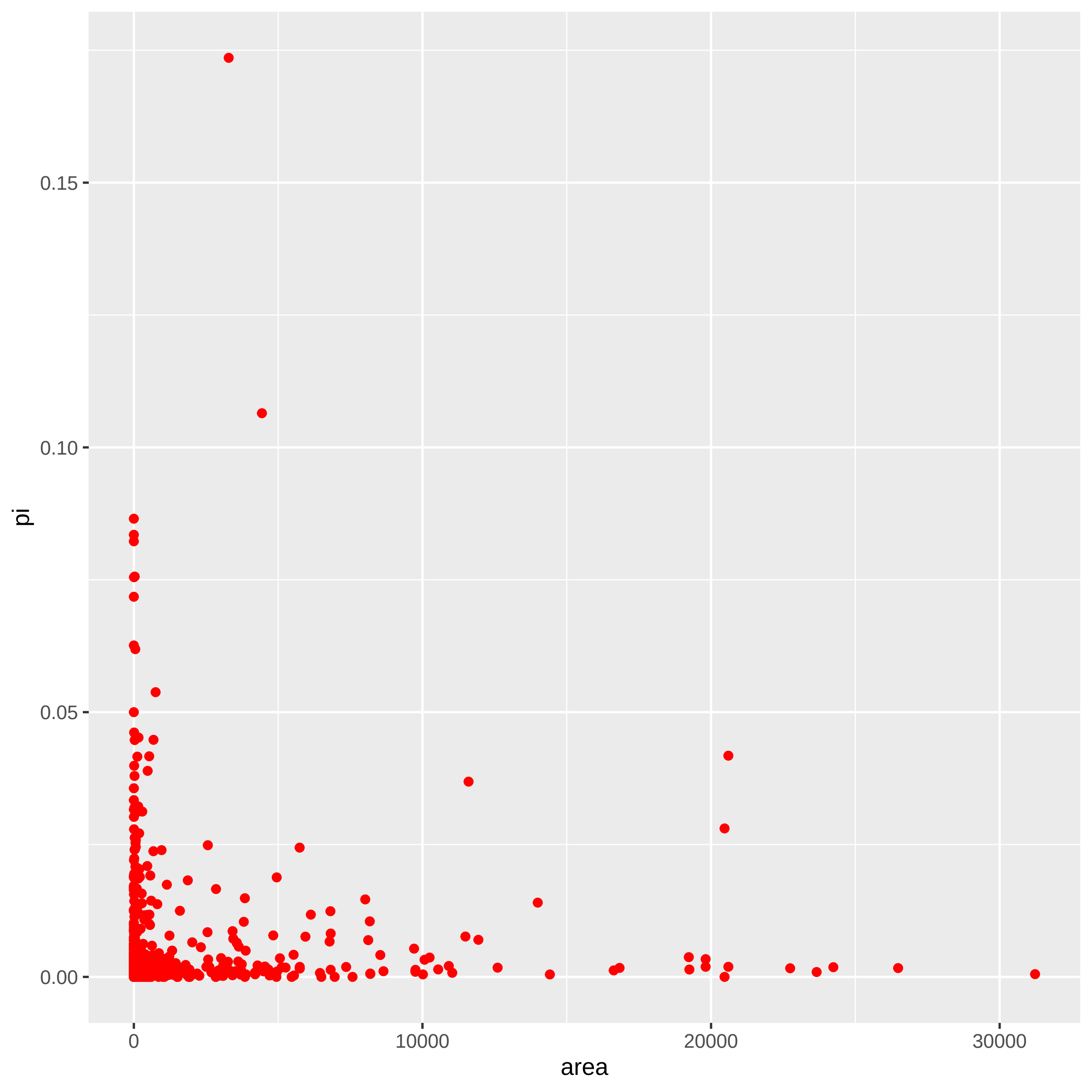

### basidiomycota_plot.pdf

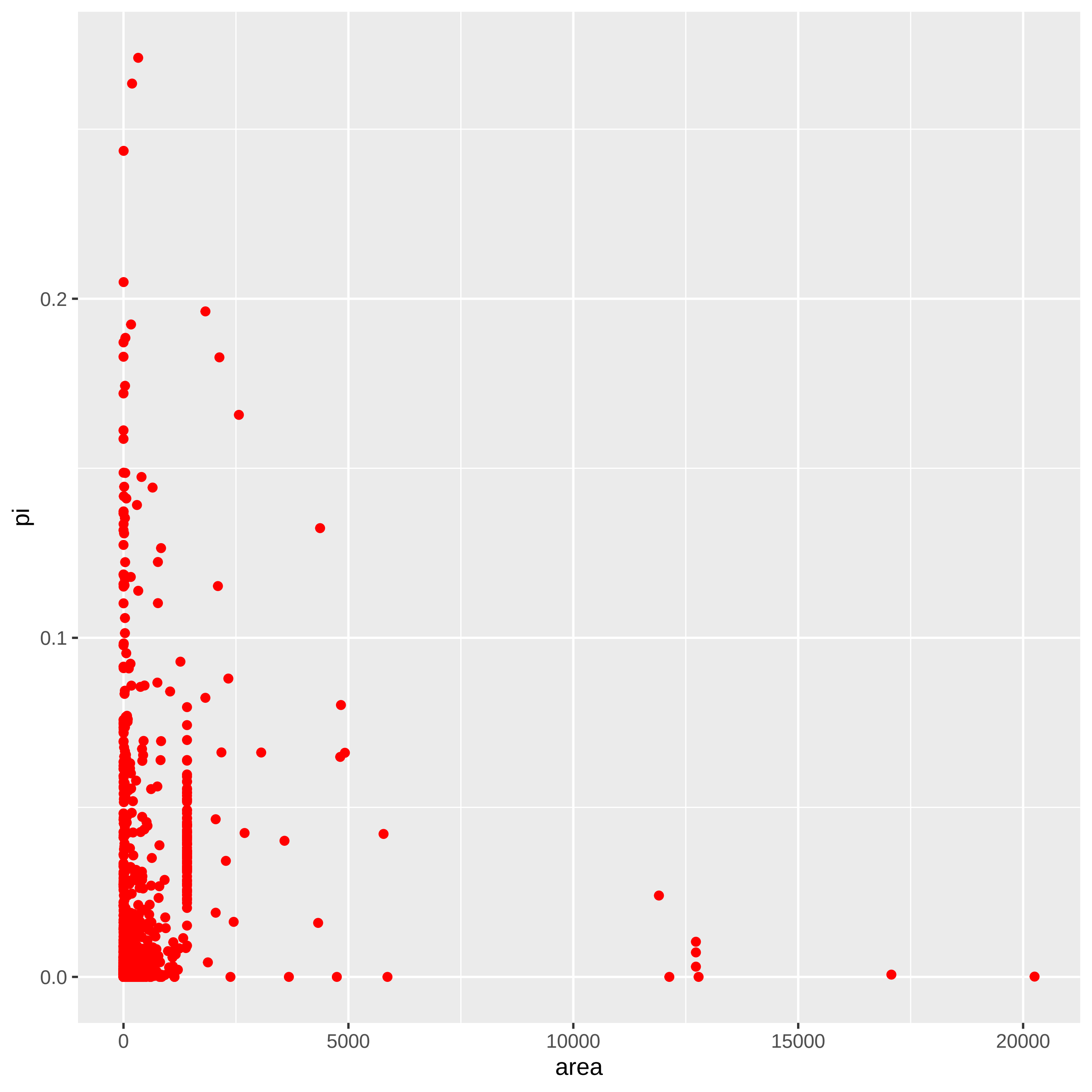

### bivalvia_plot.pdf

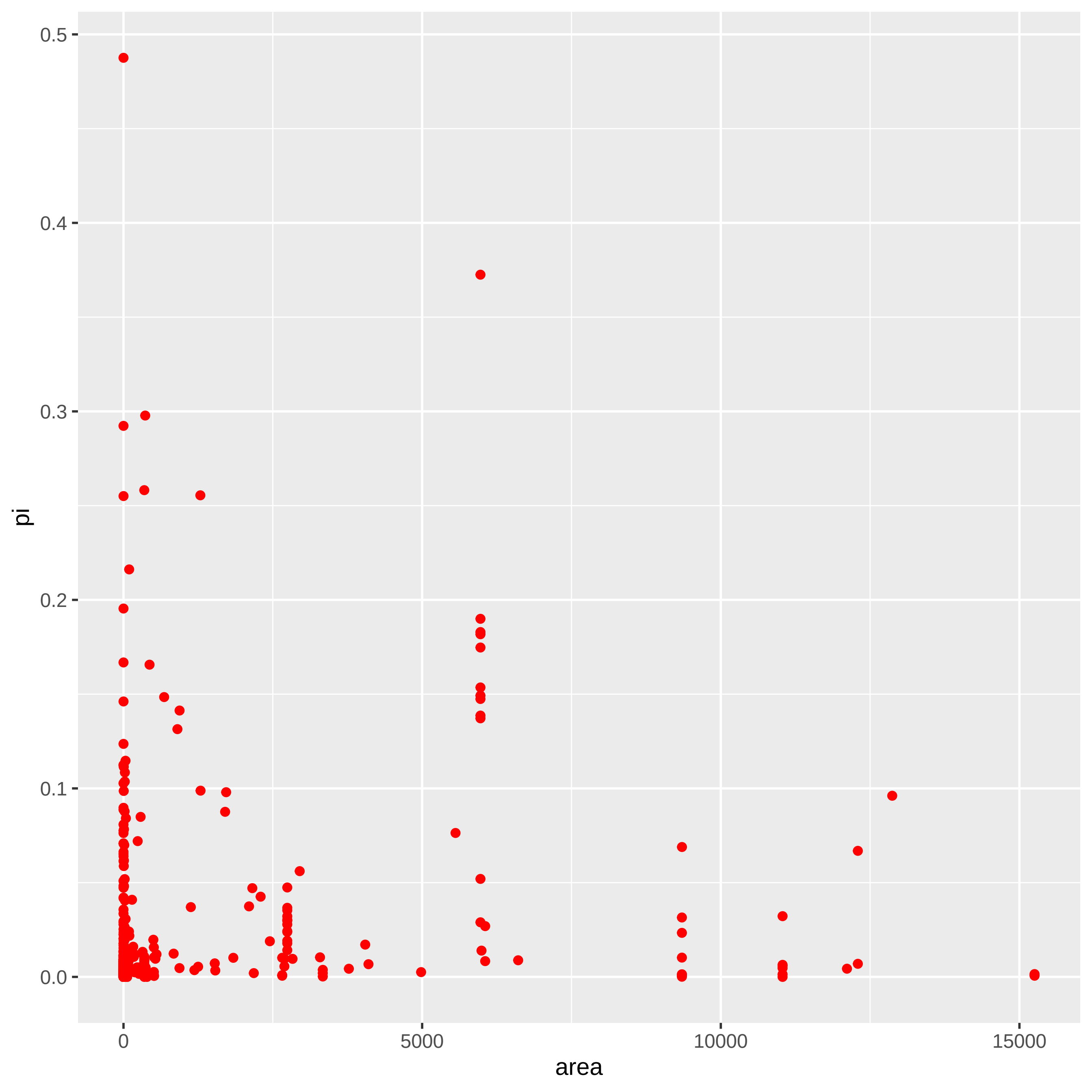

### bryophyta_plot.pdf

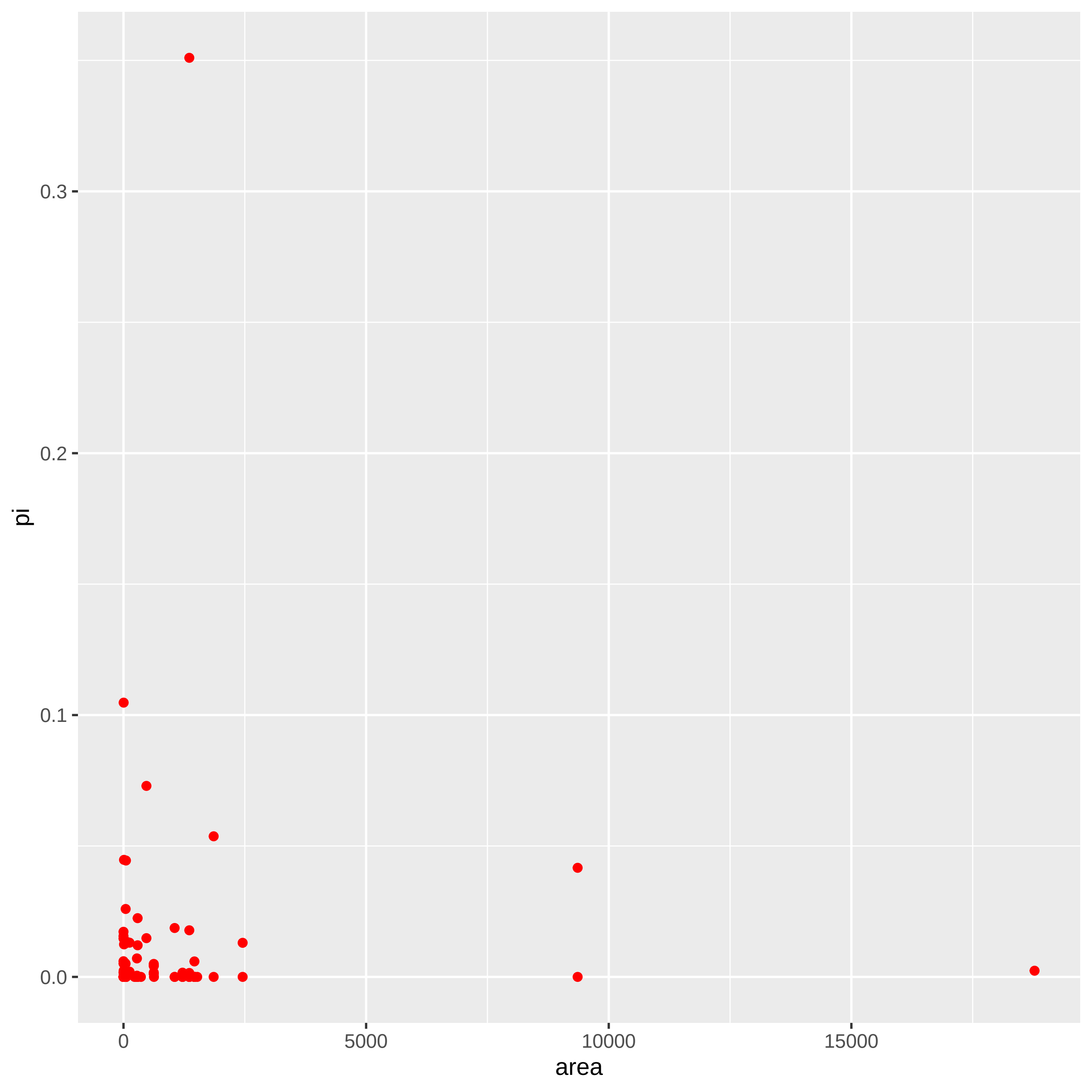

### cephalopoda_plot.pdf

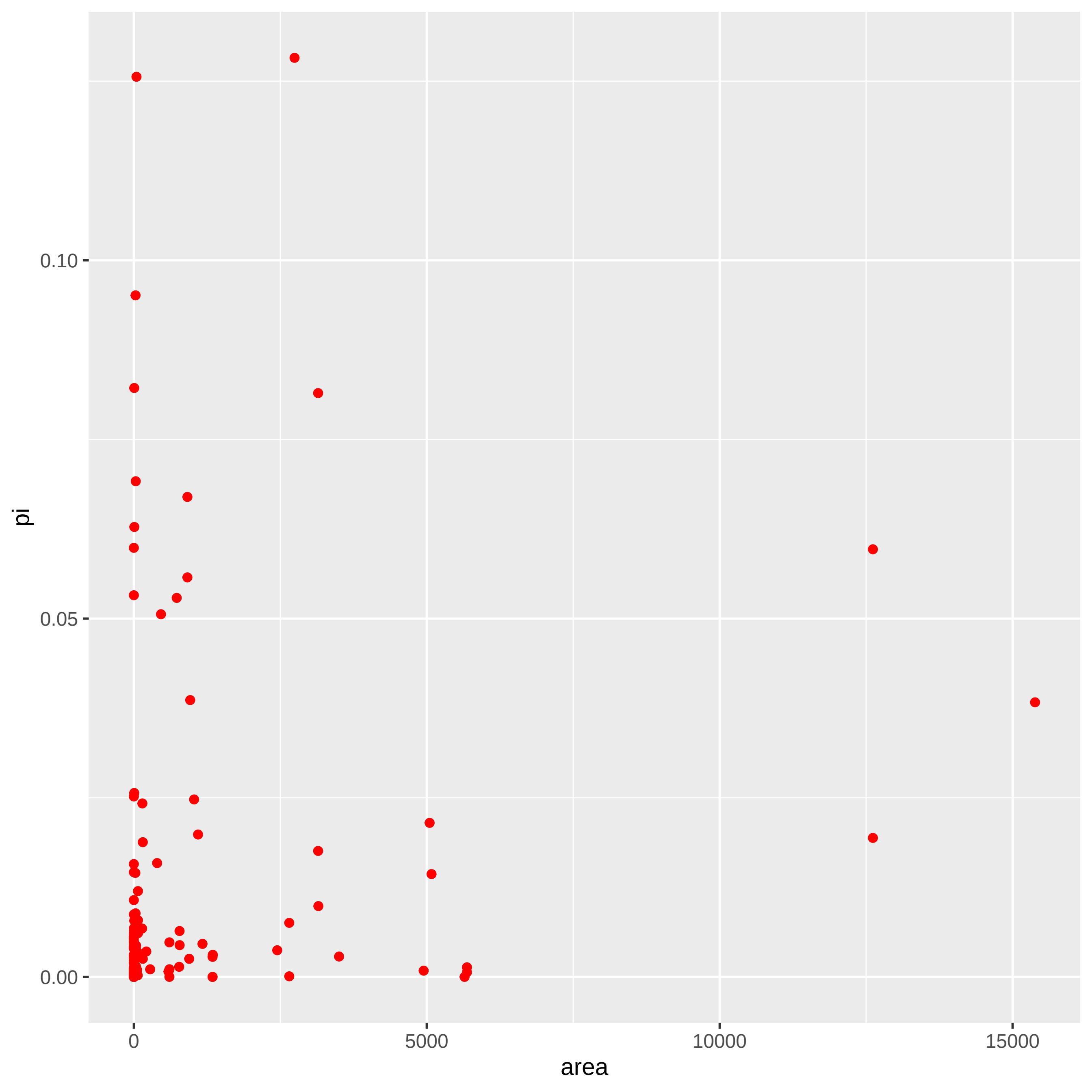

### chlorophyta_plot.pdf

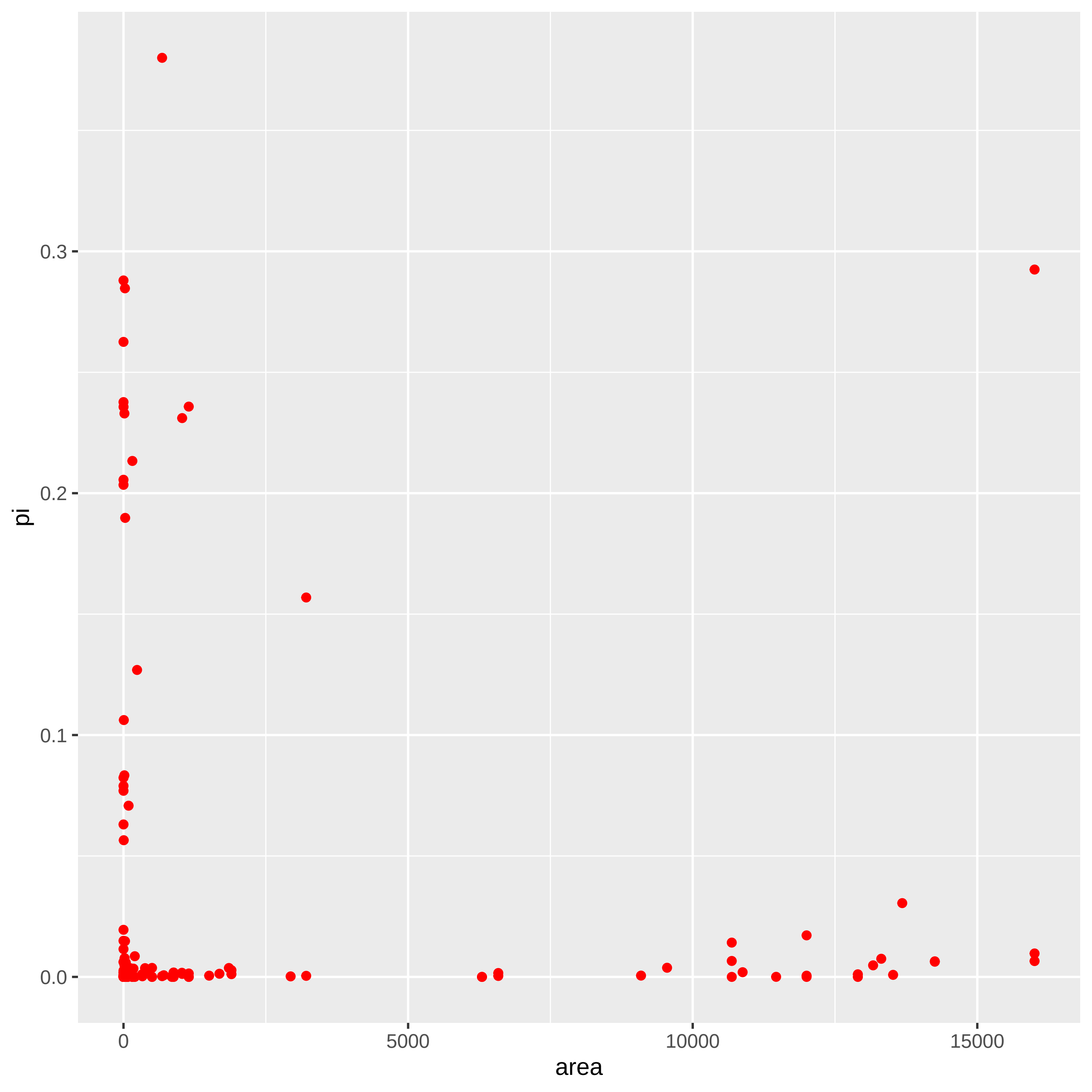

### cnidaria_plot.pdf

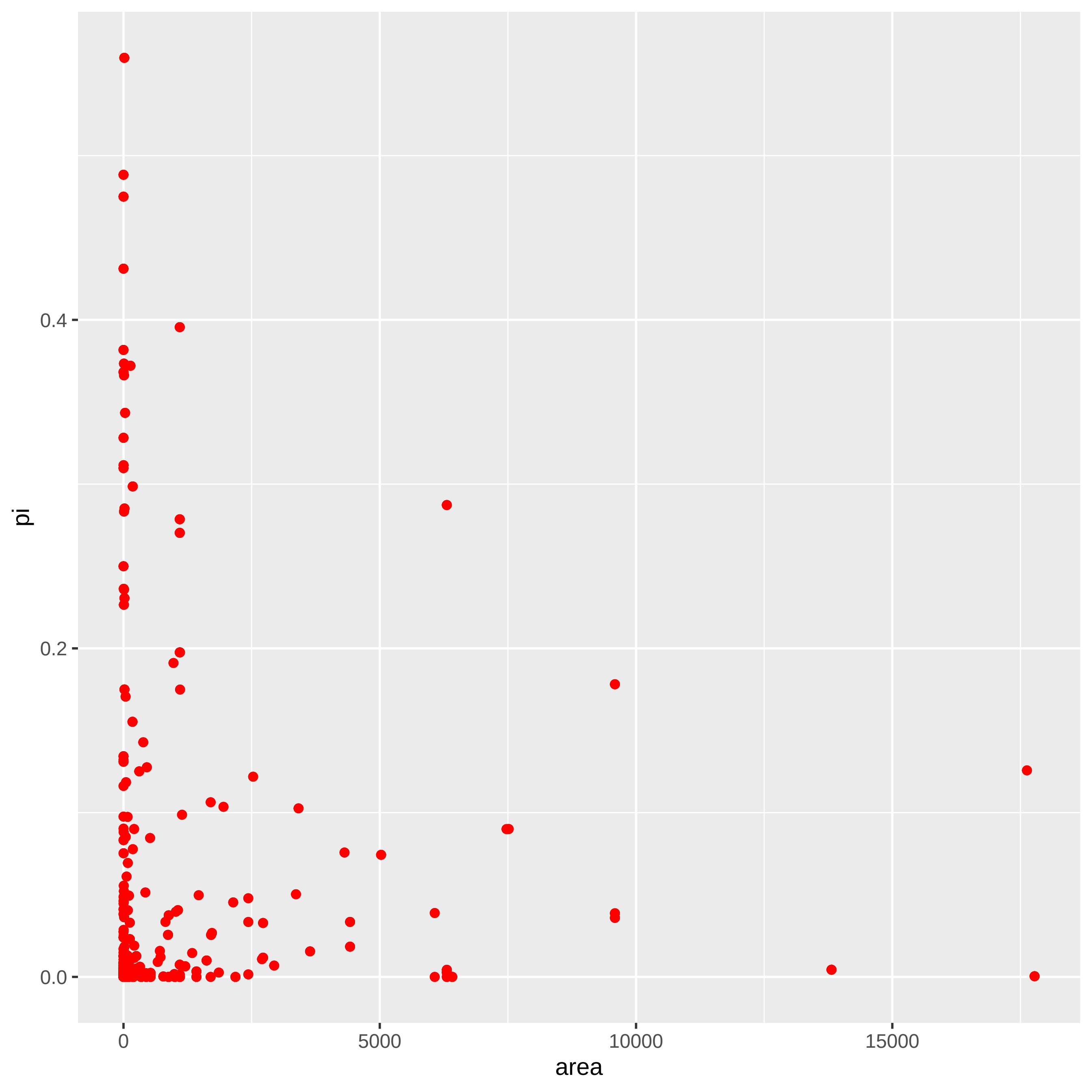

### coleoptera_plot.pdf

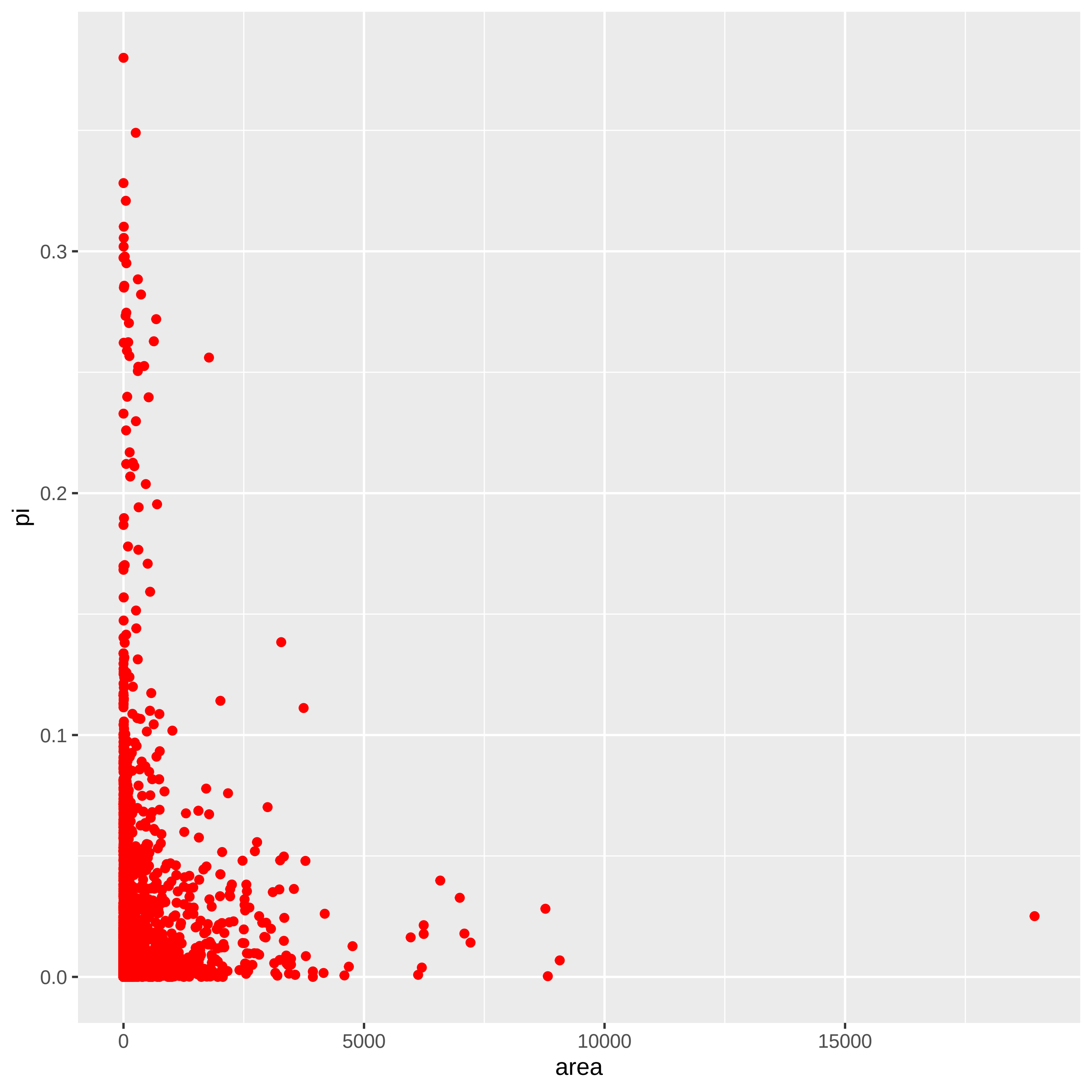

### diptera_plot.pdf

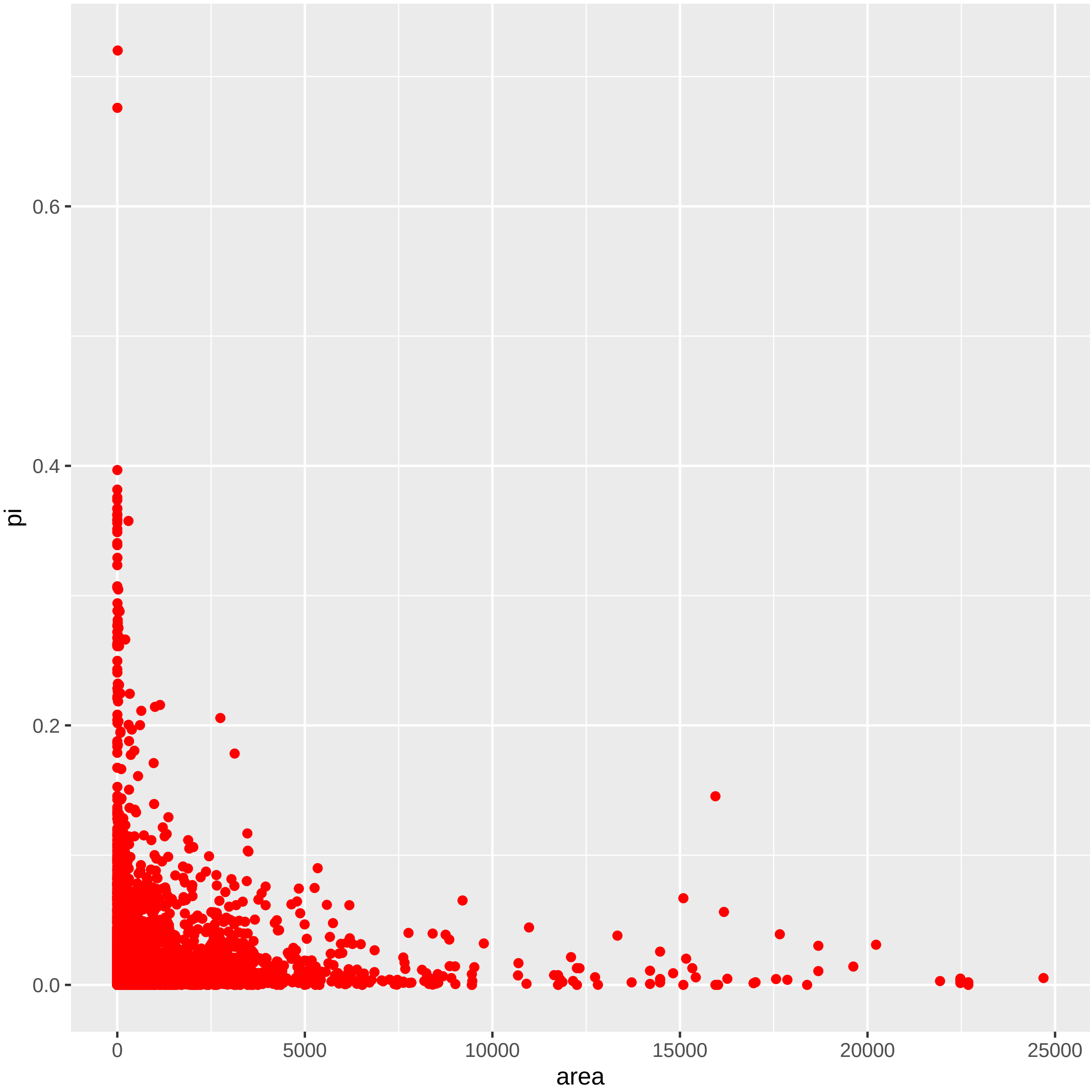

### elasmobranchii_plot.pdf

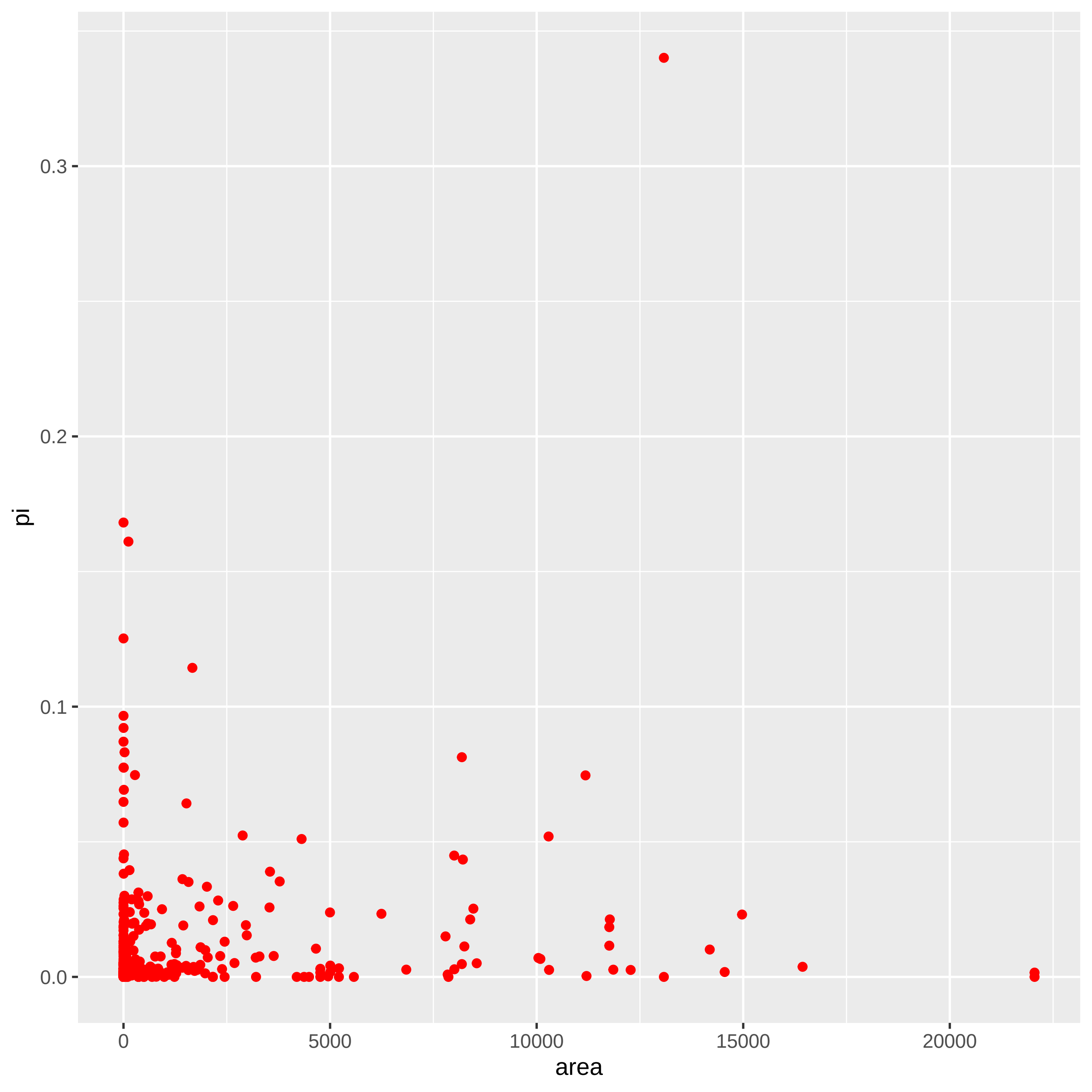

### gastropoda_plot.pdf

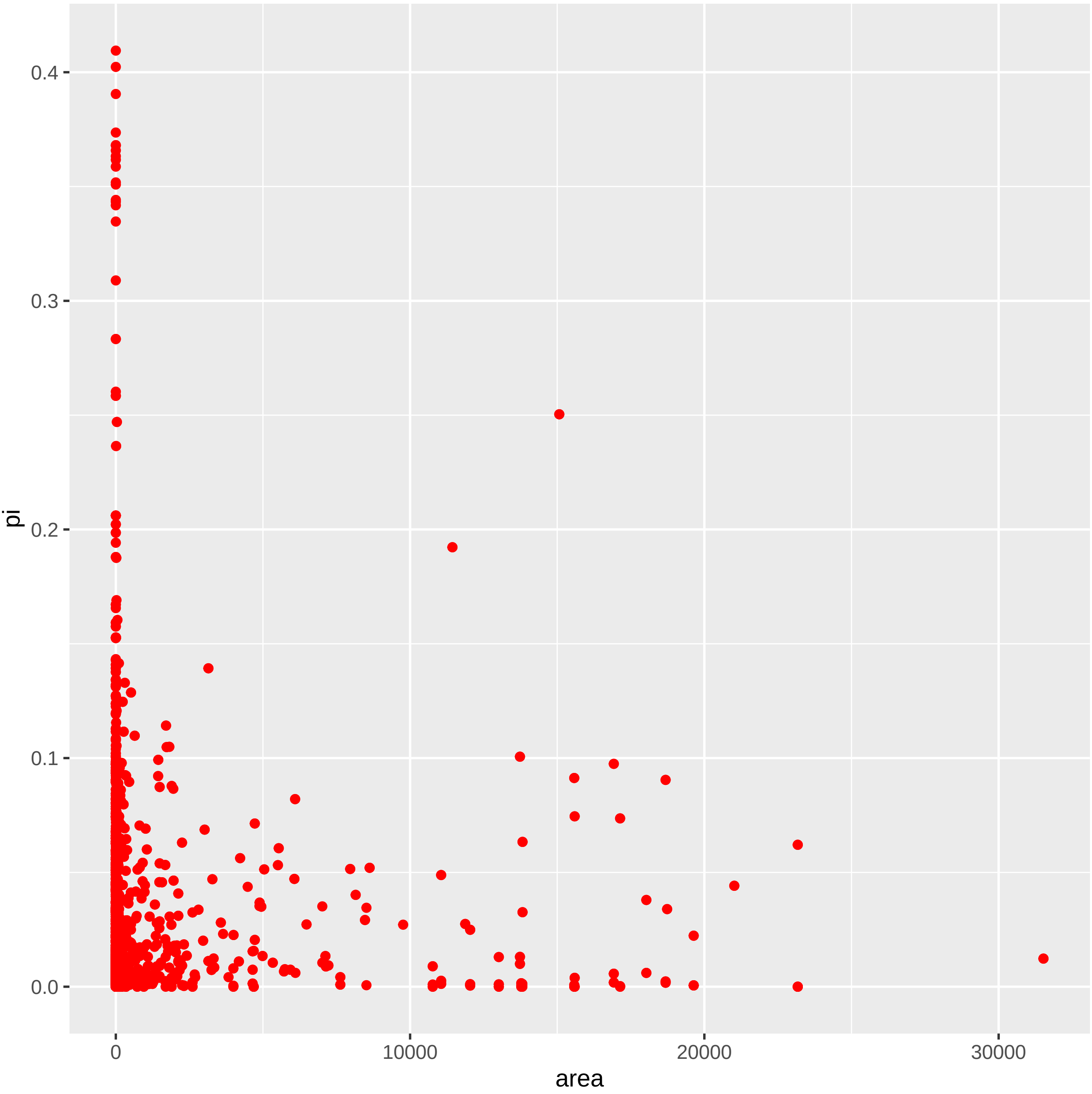

### hymenoptera_plot.pdf

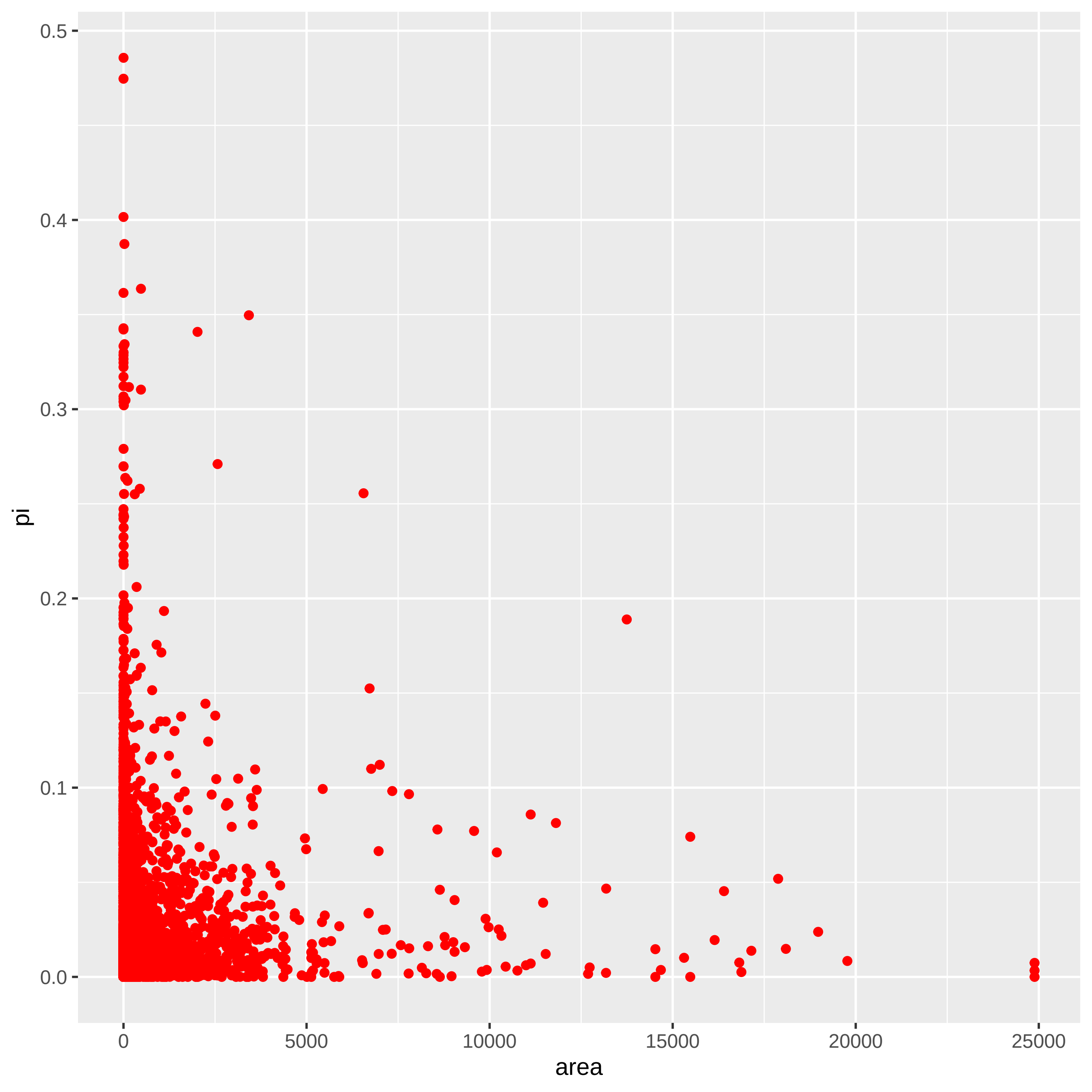

### lepidoptera_plot.pdf

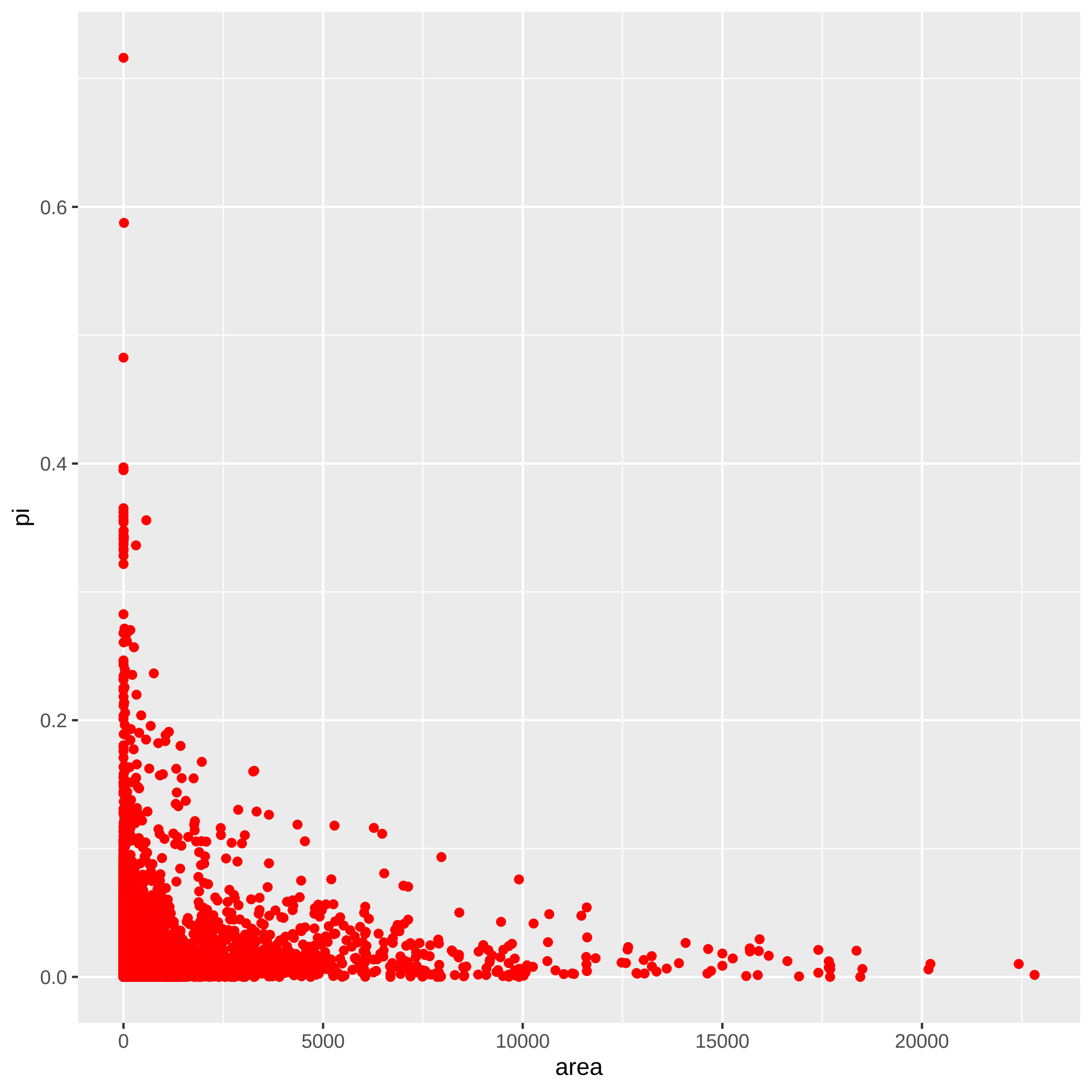

### lycopodiophyta_plot.pdf

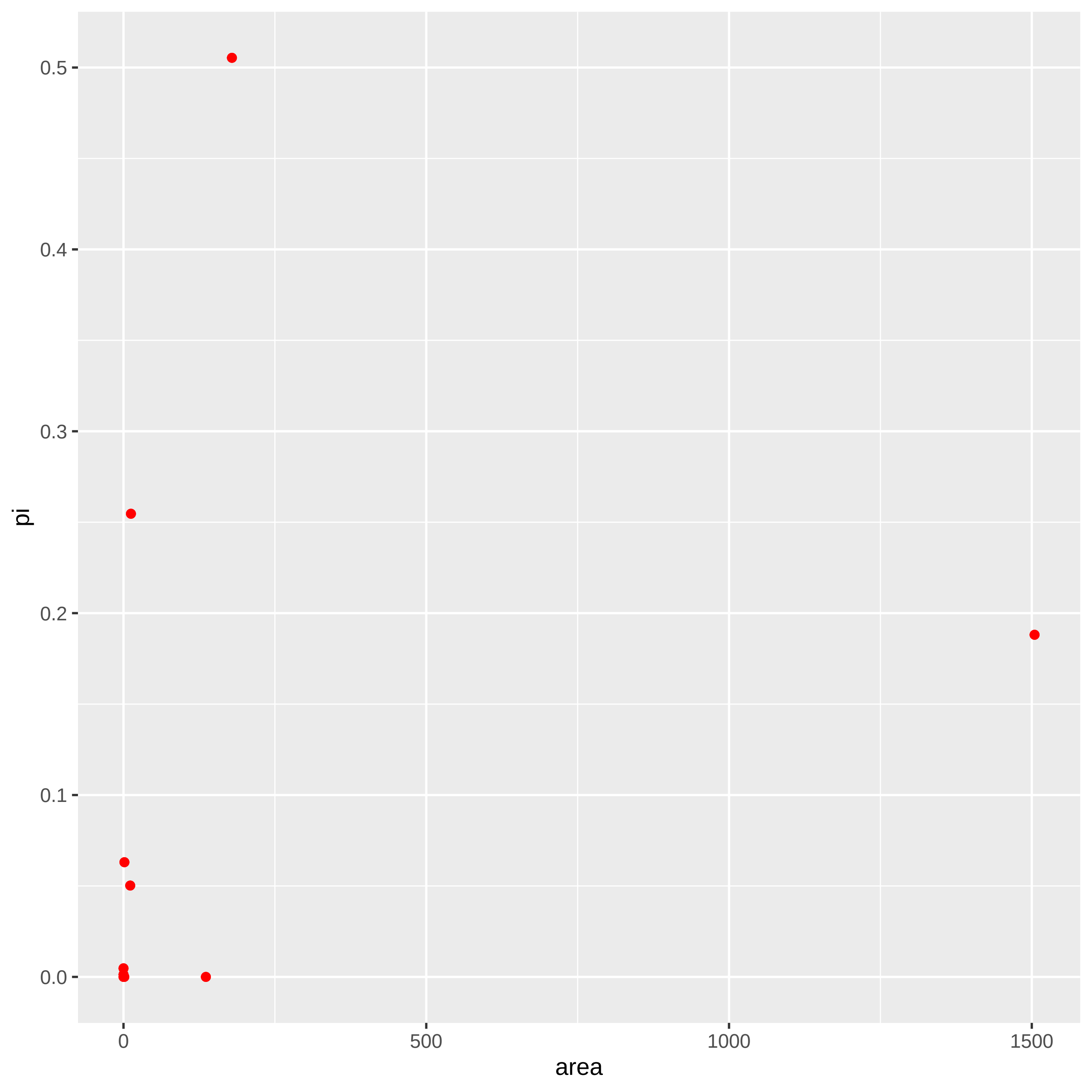

### magnoliophyta_plot.pdf

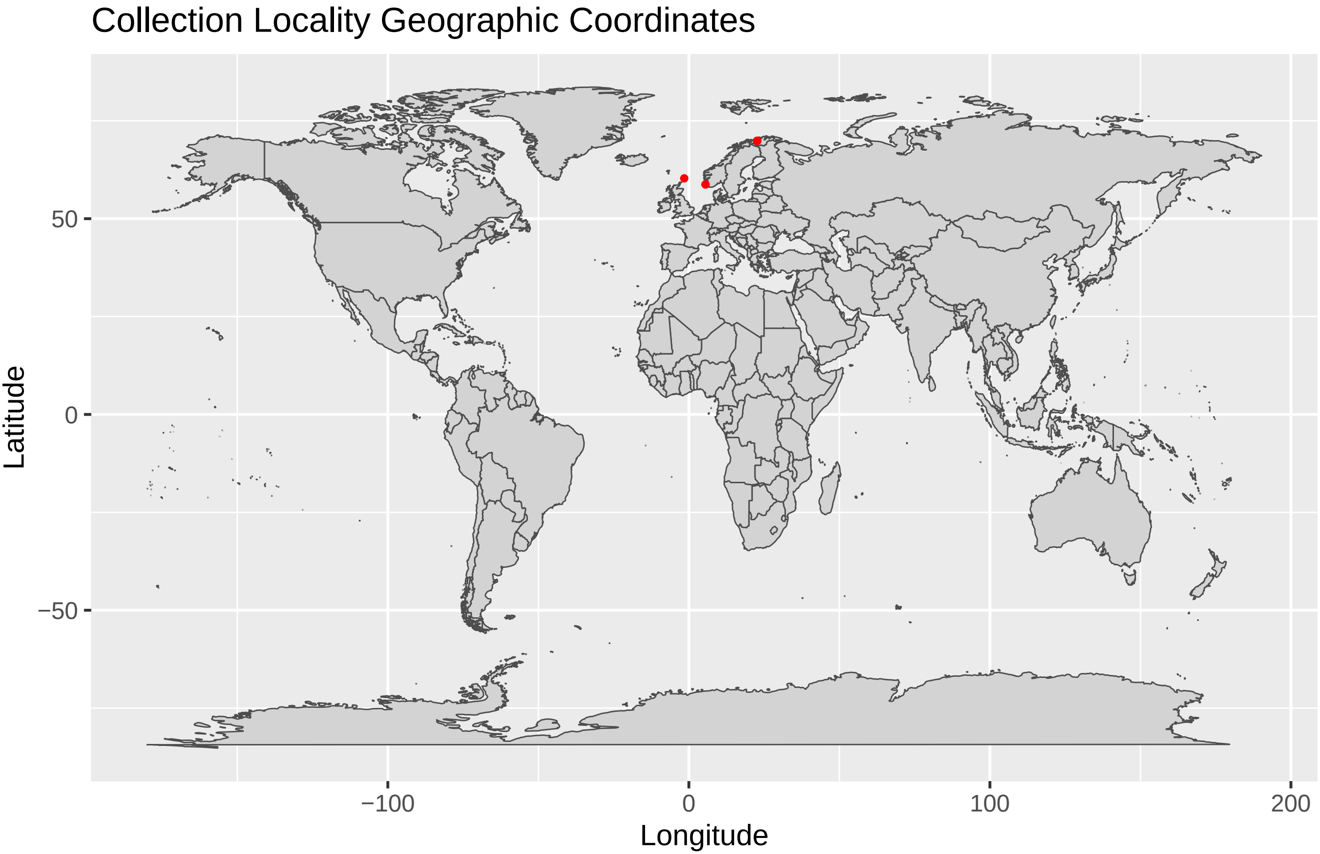

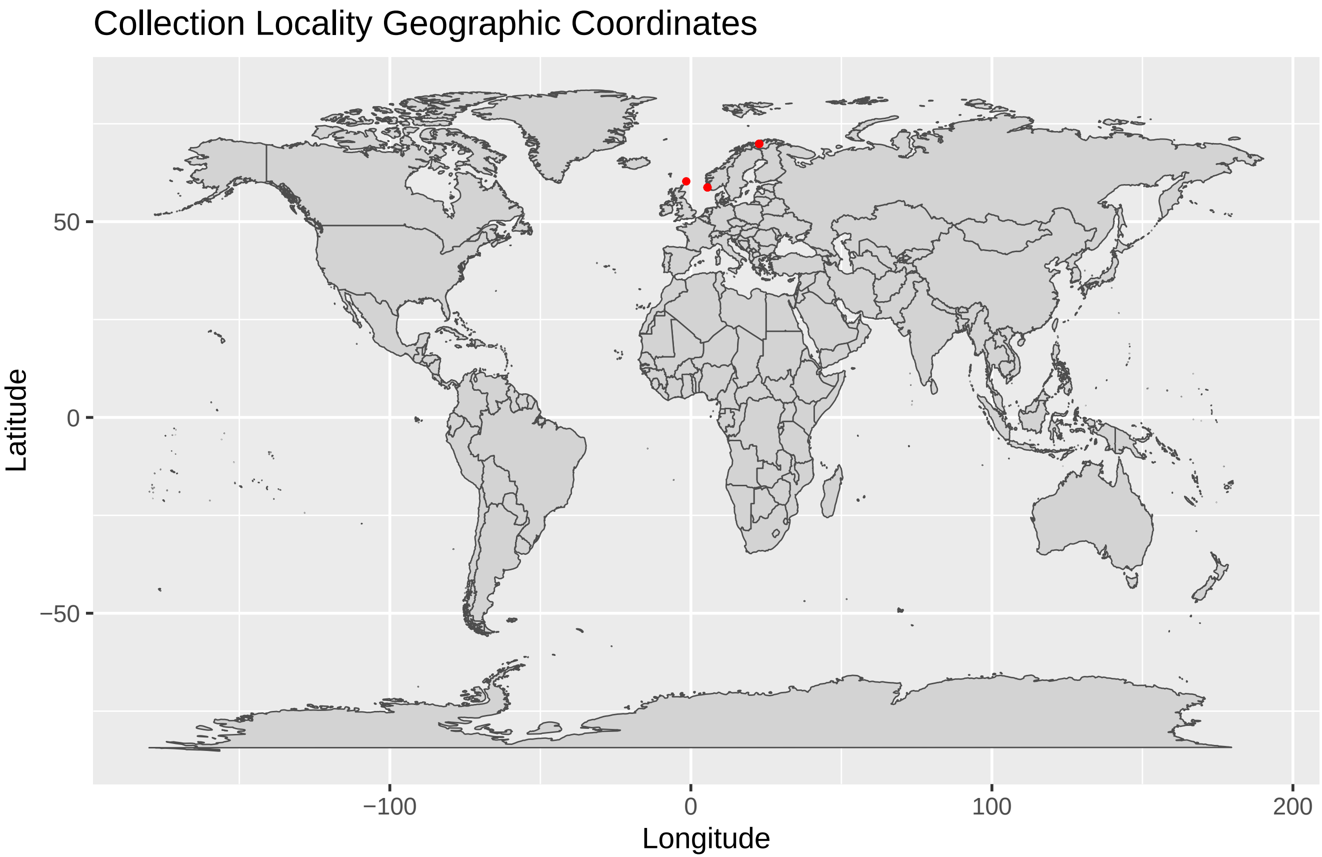

### malacostraca_plot.pdf

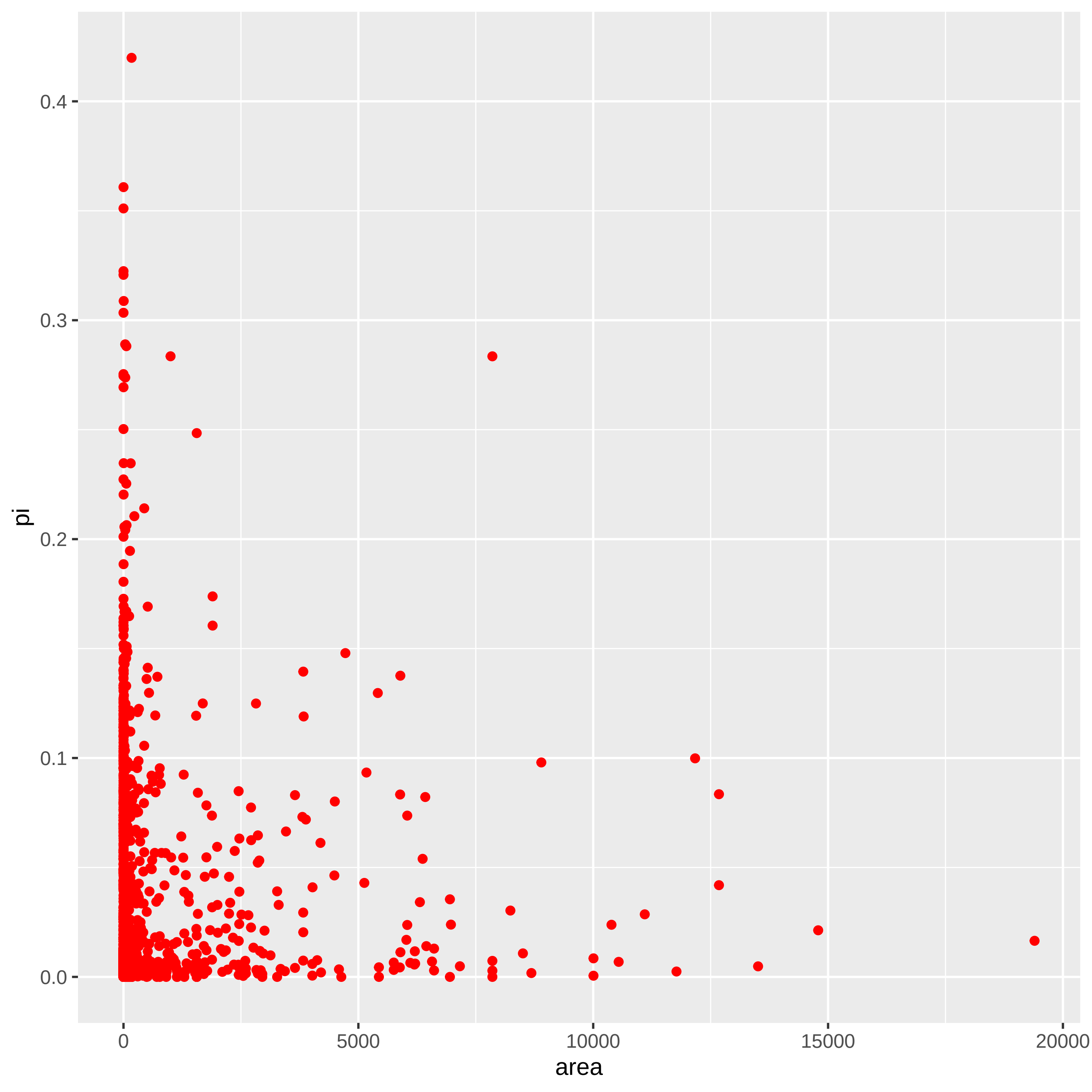

### mammalia_plot.pdf

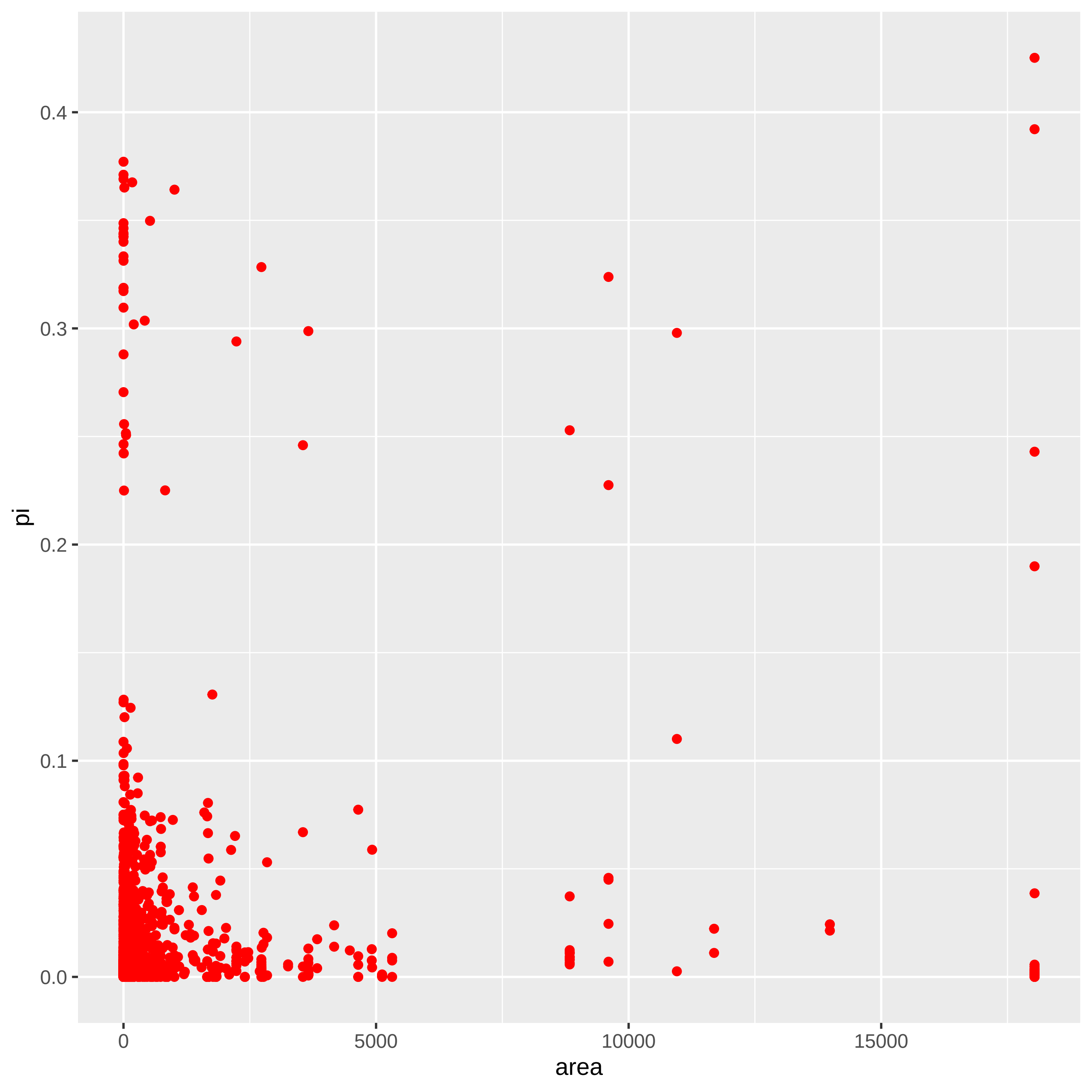

### nematoda_plot.pdf

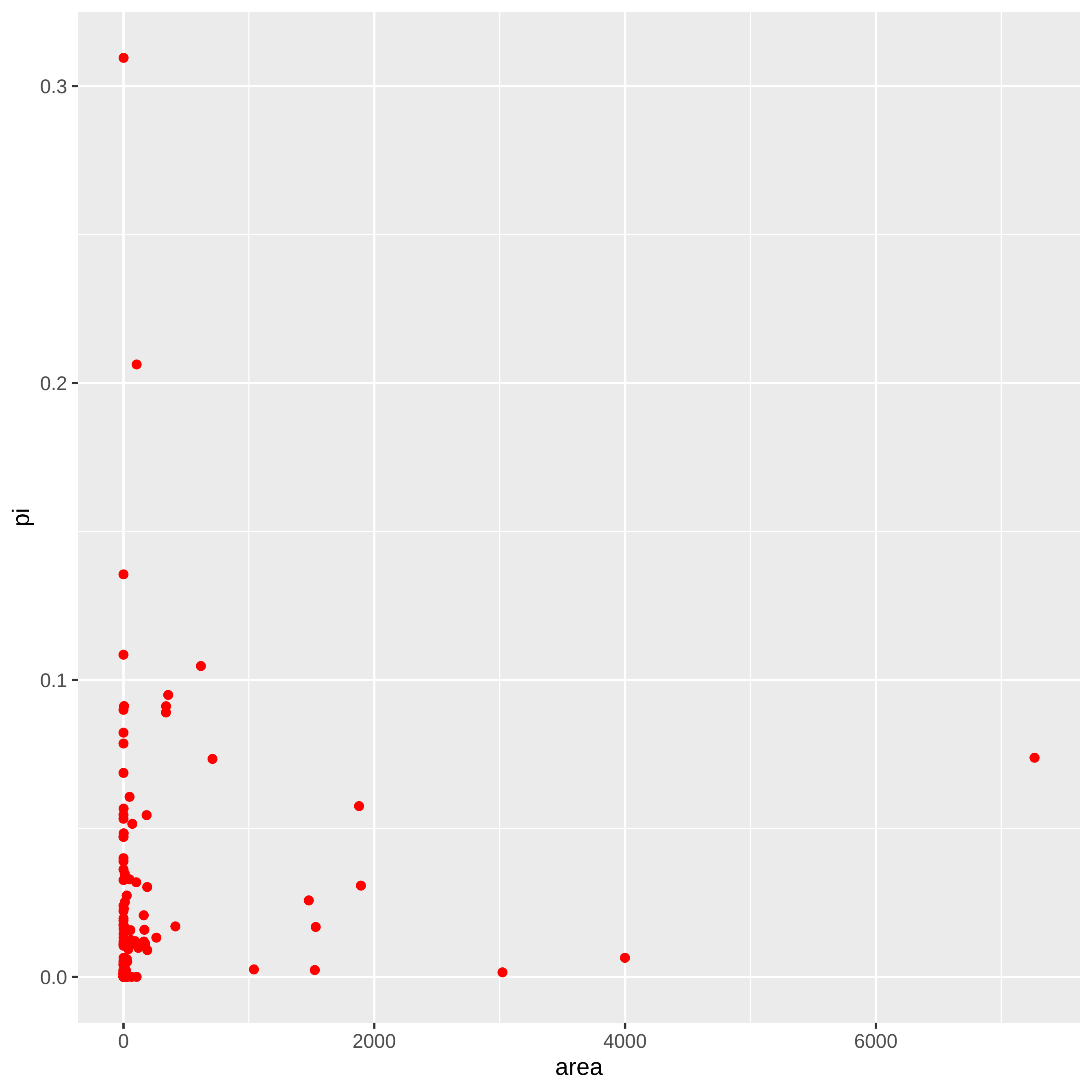

### odonata_plot.pdf

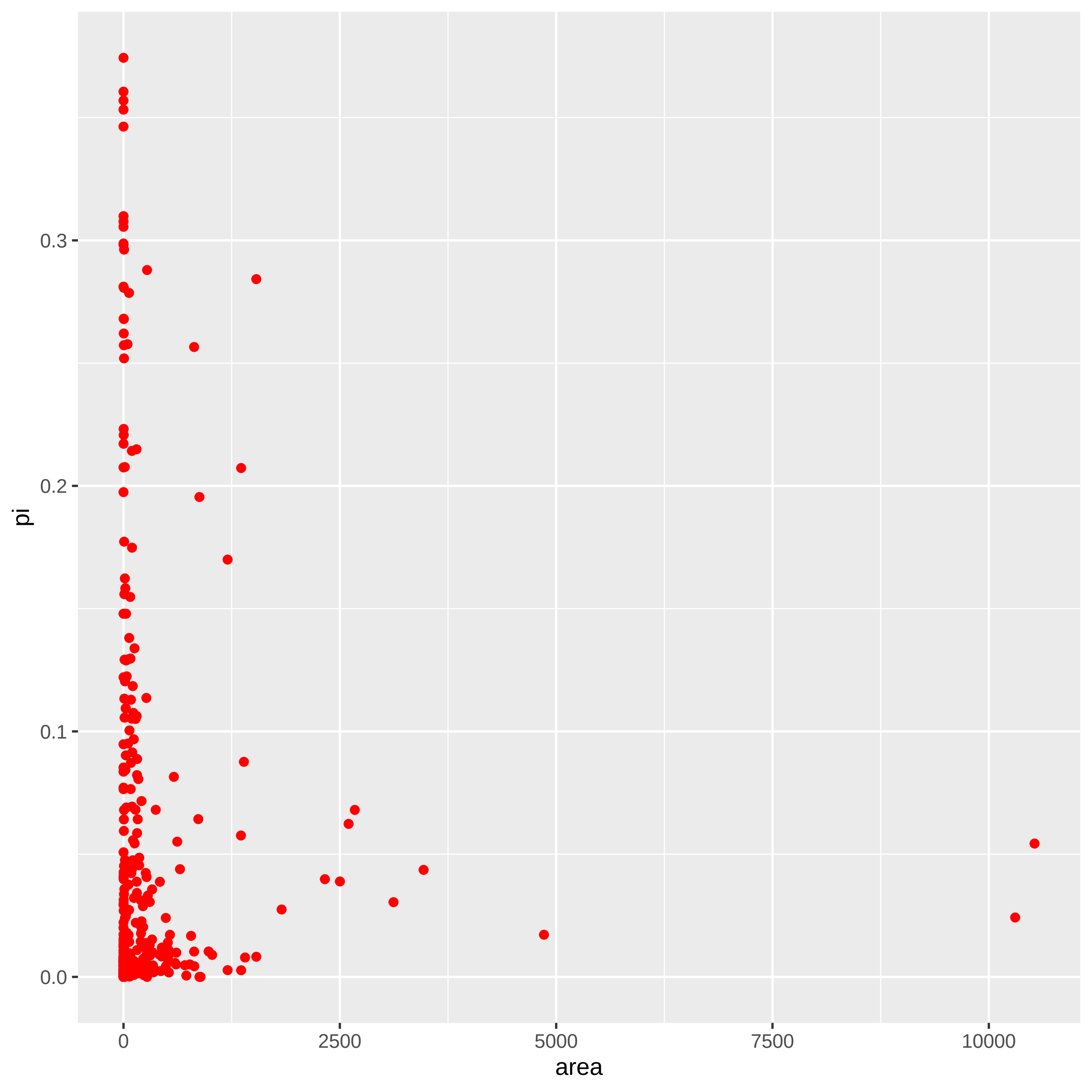

### orthoptera_plot.pdf

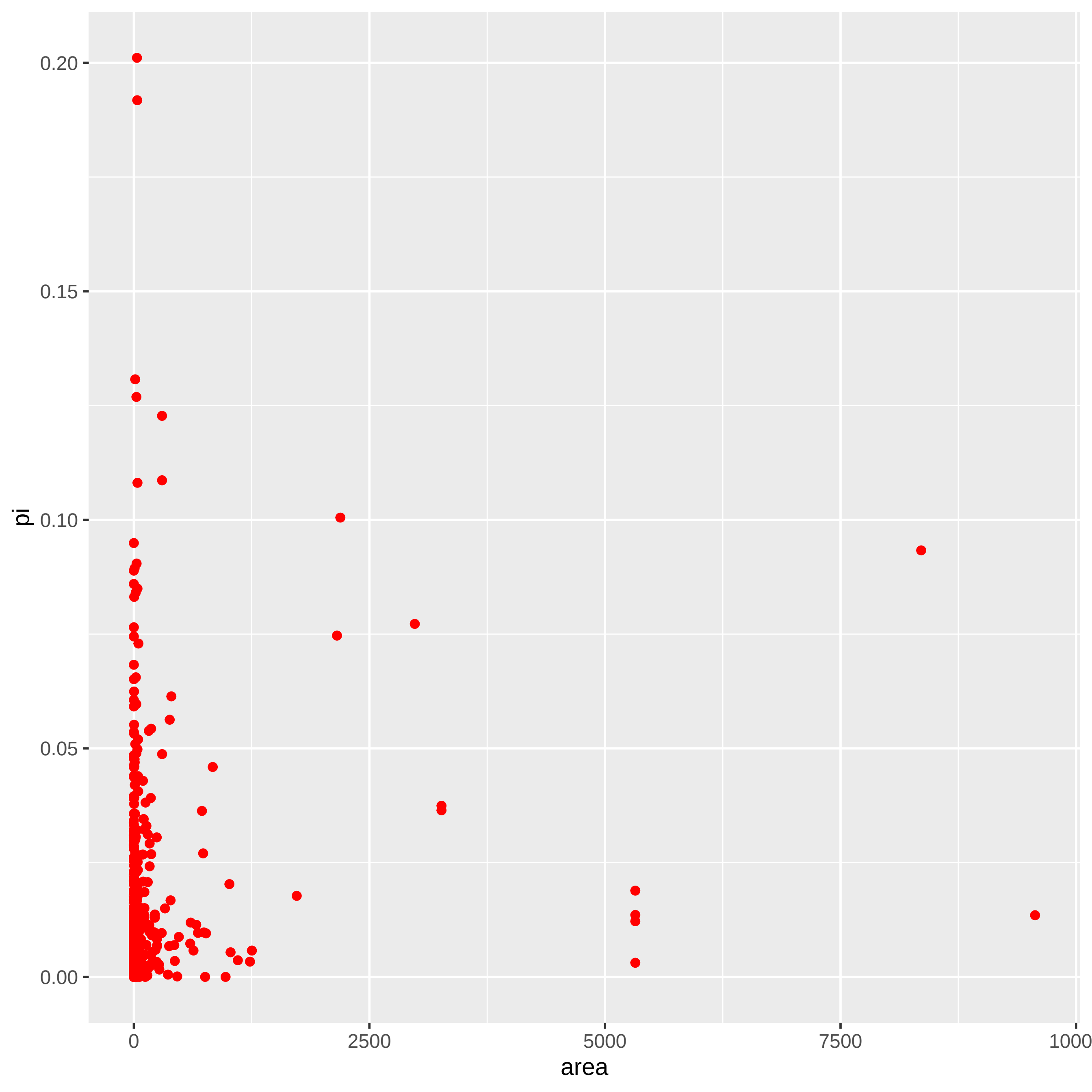

### pinophyta_plot.pdf

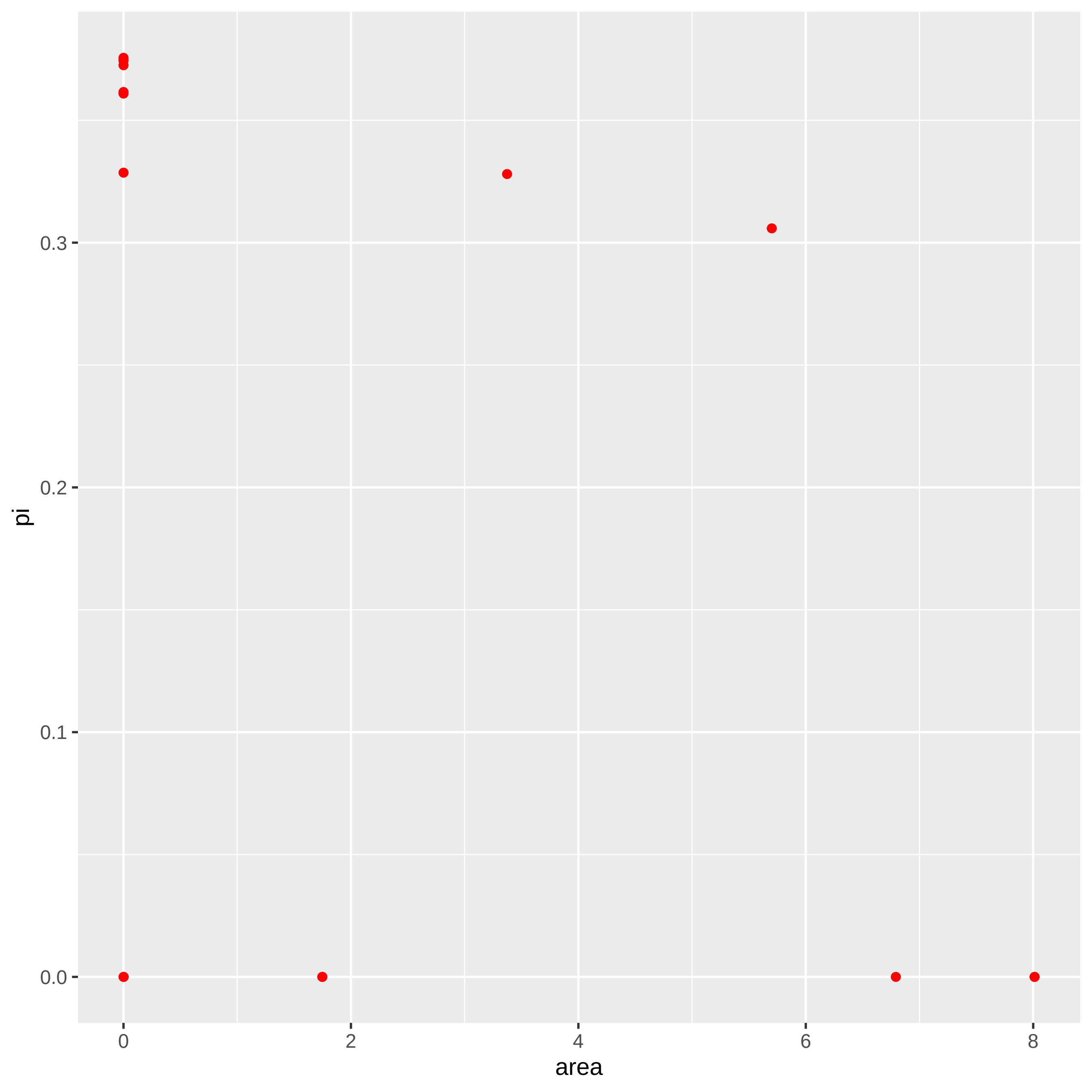

### platyhelmenthes_plot.pdf

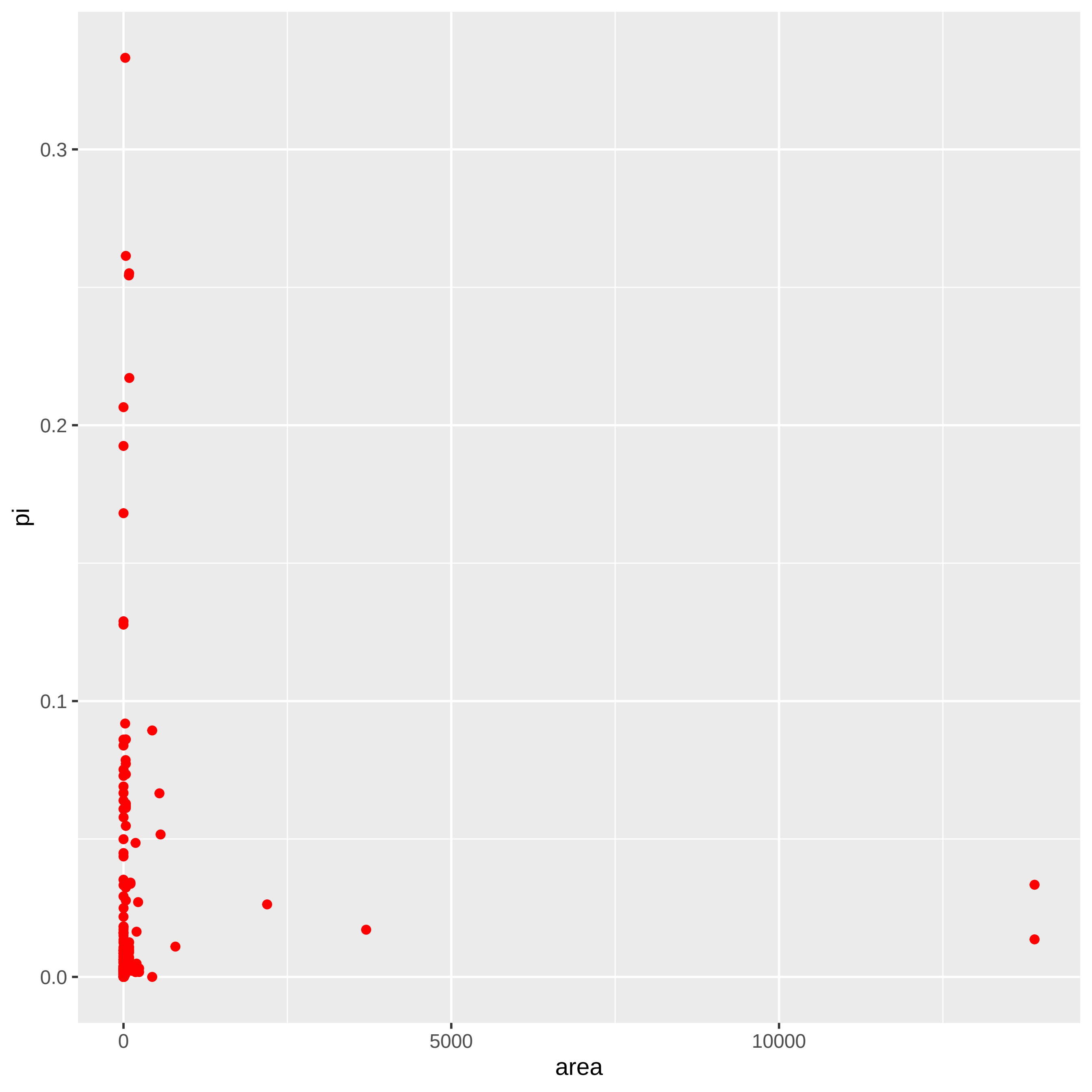

### porifera_plot.pdf

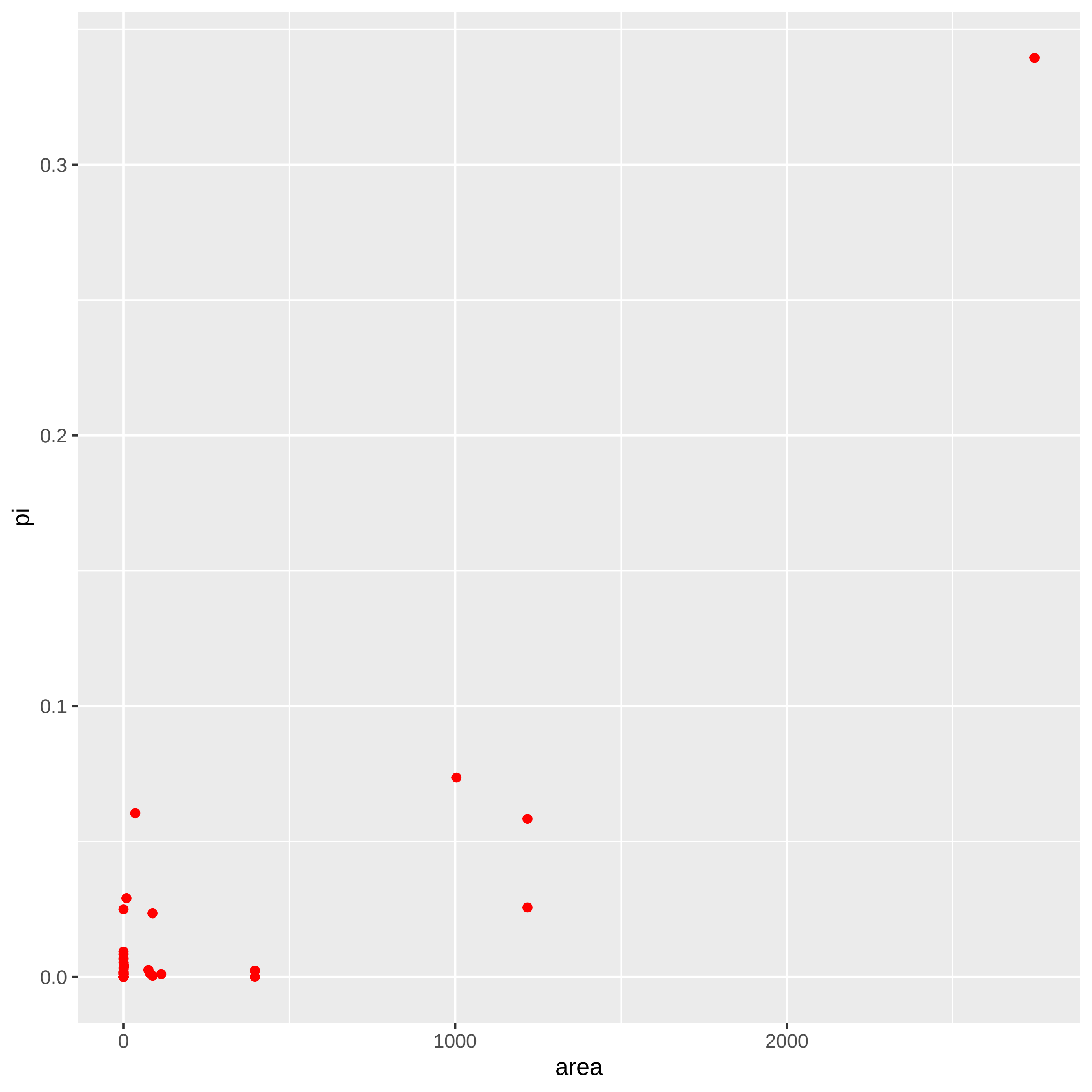

### reptilia_plot.pdf

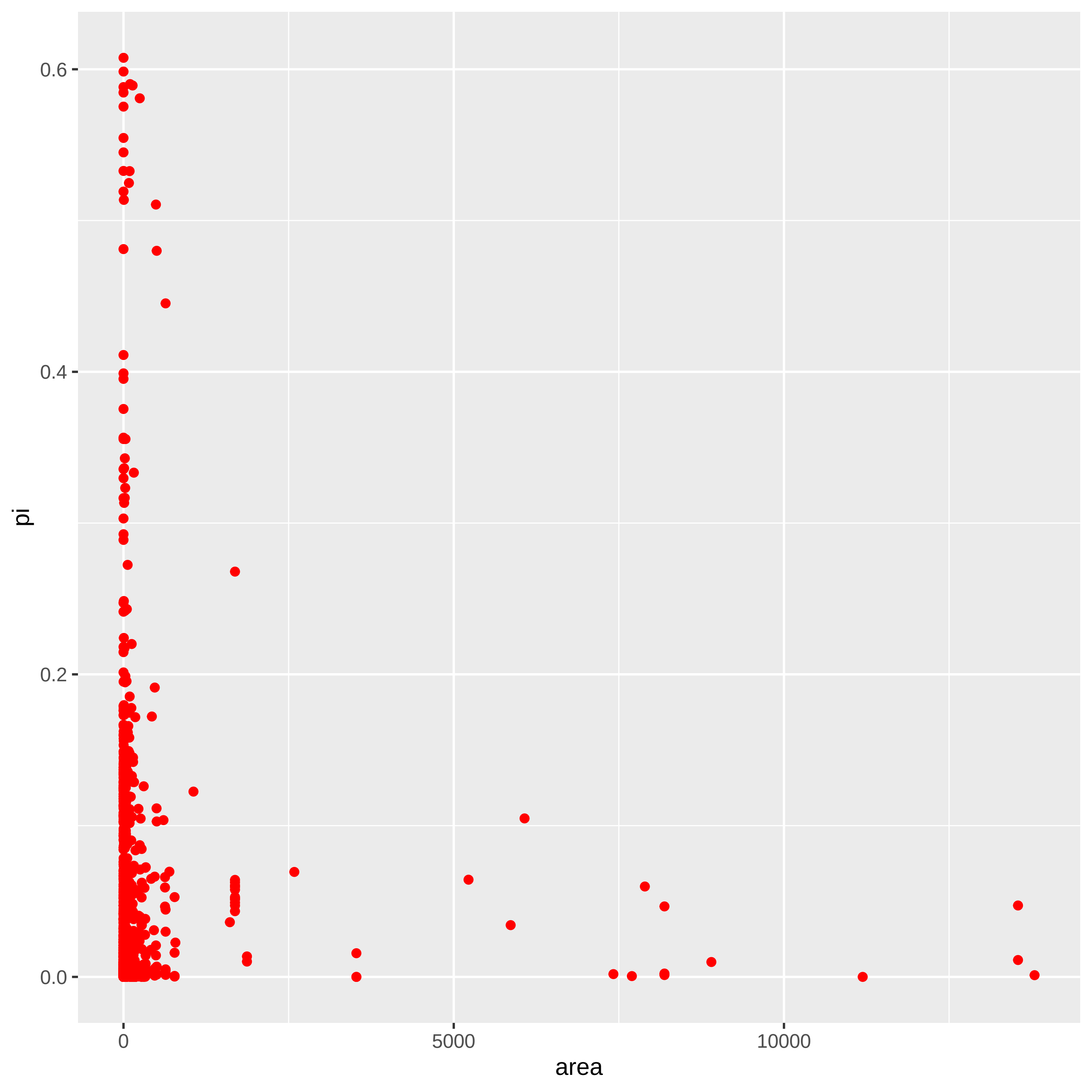

### rhodophyta_plot.pdf

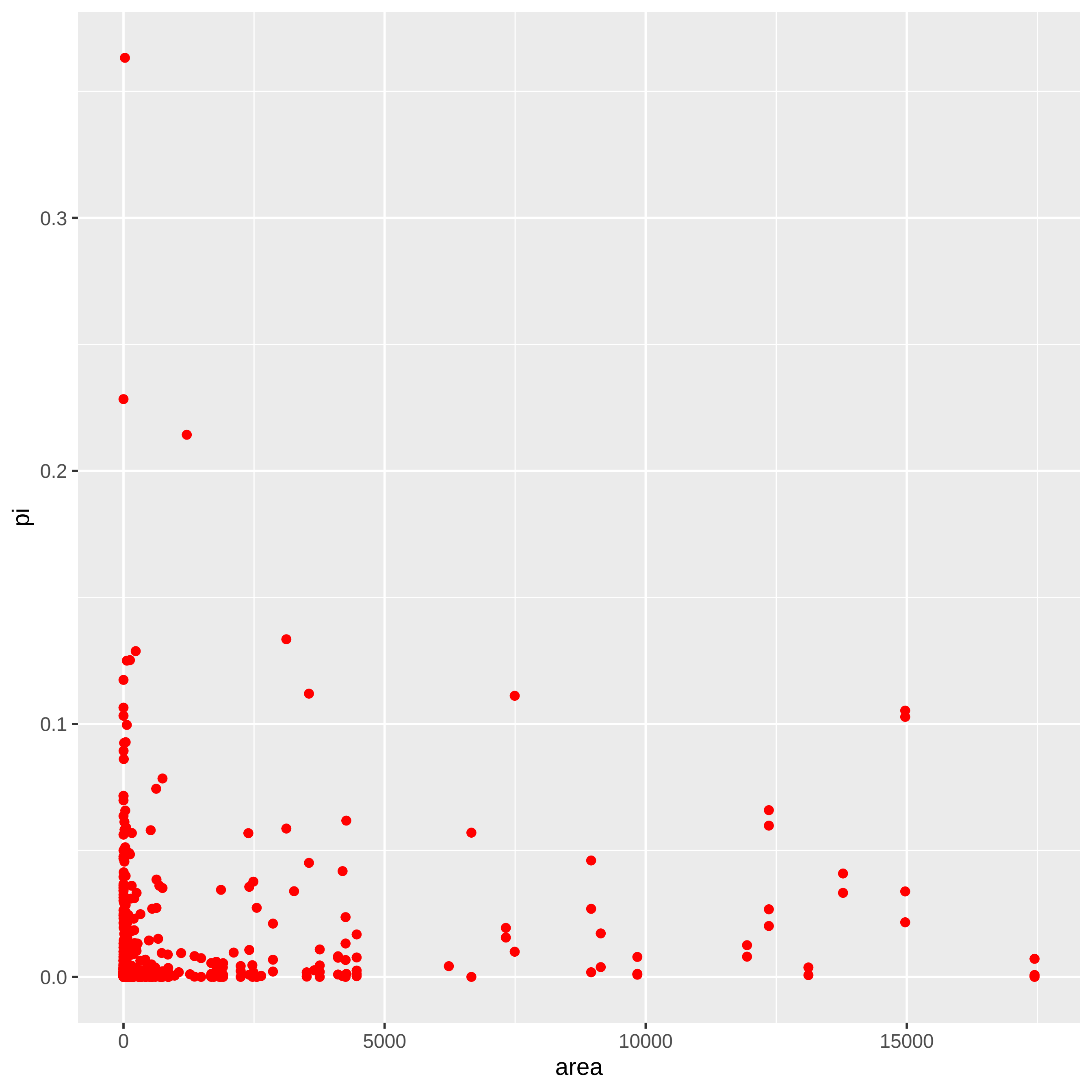

### tracheophyta_plot.pdf

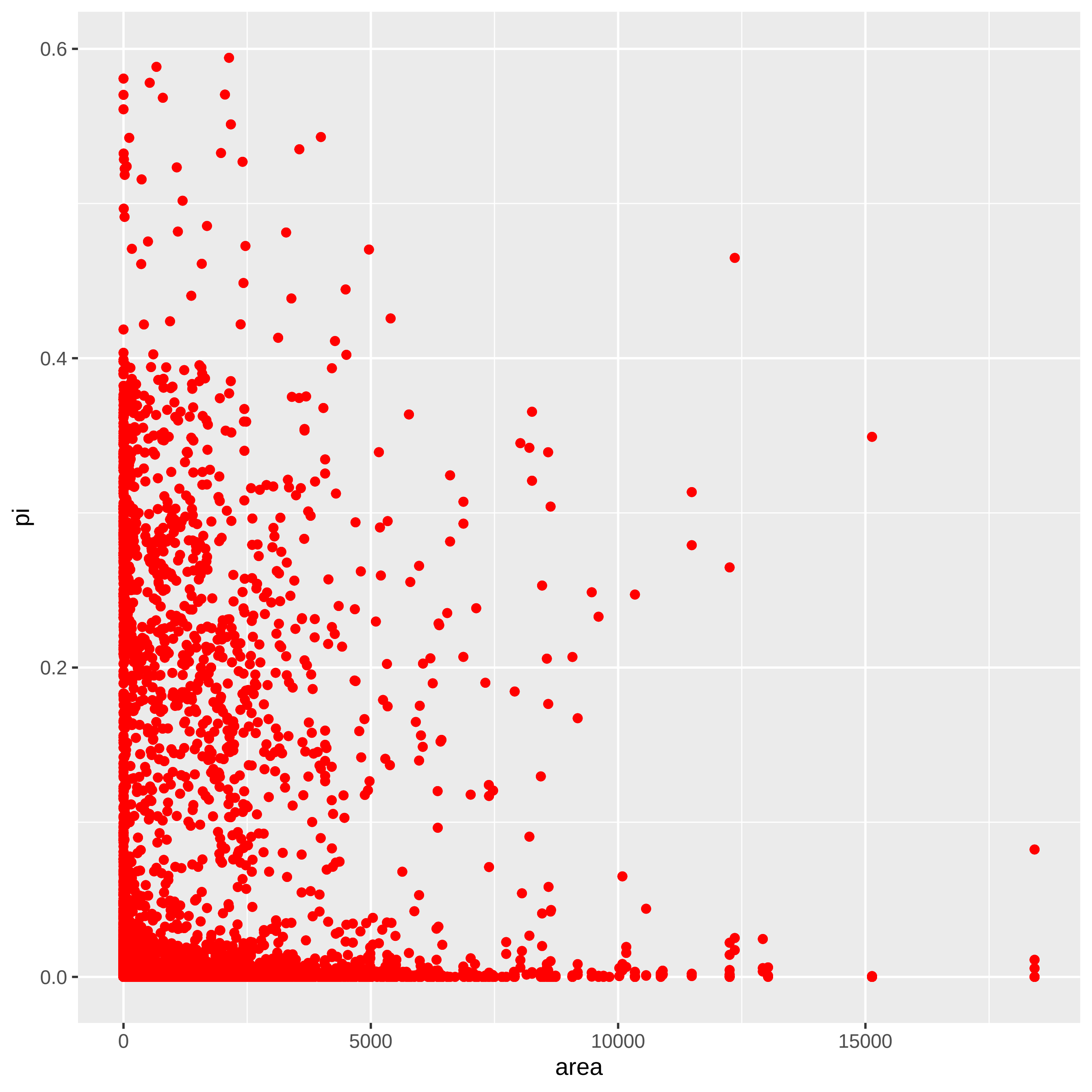
